# Identification of New *In Vivo* TonB-FepA Rendezvous Sites

**DOI:** 10.1101/2021.12.08.471779

**Authors:** Kathleen Postle, Kelvin Kho, Michael Gresock, Joydeep Ghosh, Ray Larsen

**Affiliations:** Kathleen Postle, Department of Biochemistry and Molecular Biology, The Pennsylvania State University, University Park, PA 16802, USA; School of Molecular Biosciences, Washington State University, Pullman, WA 99163; Institut Pasteur, Unité Biologie et Génétique de la Paroi Bactérienne, Département de Microbiologie, Paris 75015, France; Department of Biology, University of Mt. Union, Alliance OH, 44601; Division of Cellular and Gene Therapies, US FDA Center for Biologics Evaluation and Research, Silver Spring, MD; Department of Biological Sciences, Bowling Green State University, Bowling Green OH, 43403

**Keywords:** TonB, FepA, disulfide bond, amphipathic helix, energy transduction

## Abstract

The TonB system of Gram-negative bacteria uses the protonmotive force of the cytoplasmic membrane to energize active transport of large or scarce nutrients across the outer membrane by means of customized beta-barrels known as TonB-dependent transporters (TBDTs). The lumen of each TBDT is occluded by an amino-terminal domain, called the cork, which must be displaced for transport of nutrients or translocation of the large protein toxins that parasitize the system. A complex of cytoplasmic membrane proteins consisting of TonB, ExbB and ExbD harnesses the protonmotive force that TonB transmits to the TBDT. The specifics of this energy transformation are a source of continuing interest. The amino terminal domain of a TBDT contains a region called the TonB box, that is essential for the reception of energy from TonB. This domain is the only identified site of *in vivo* interaction between the TBDT and TonB, occurring through a non-essential region centered on TonB residue Q160. Because TonB binds to TBDTs whether or not it is active or even intact, the mechanism and extent of cork movement *in vivo* has been challenging to discover. In this study, we used *in vivo* disulfide crosslinking between eight engineered Cys residues in *Escherichia coli* TonB and 42 Cys substitutions in the TBDT FepA, including the TonB box, to identify novel sites of interaction *in vivo*. The TonB Cys substitutions in the core of an essential carboxy terminal amphipathic helix (residues 199-216) were compared to TonB Q160C interactions. Functionality of the *in vivo* interactions was established when the presence of the inactive TonB H20A mutation inhibited them. A previously unknown functional interaction between the hydrophilic face of the amphipathic helix and the FepA TonB box was identified. Interaction of Q160C with the FepA TonB box appeared to be less functionally important. The two different parts of TonB also differed in their interactions with the FepA cork and barrel turns. While the TonB amphipathic helix Cys residues interacted only with Cys residues on the periplasmic face of the FepA cork, TonB Q160C interacted with buried Cys substitutions within the FepA cork, the first such interactions seen with any TBDT. Both sets of interactions required active TonB. Taken together, these data suggest a model where the amphipathic helix binds to the TonB box, causing the mechanically weak domain of the FepA cork to dip sufficiently into the periplasmic space for interaction with the TonB Q160 region, which is an interaction that does not occur if the TonB box is deleted. The TonB amphipathic helix also interacted with periplasmic turns between FepA β-strands *in vivo* supporting a surveillance mechanism where TonB searched for TBDTs on the periplasmic face of the outer membrane.

## INTRODUCTION

The TonB system of *Escherichia coli* appears to be an answer to the challenges posed to Gram-negative bacteria by their dual membrane cell envelope. In particular, the outer membrane is a largely protective sieve with a diffusion cut-off of around 600 Da in *E. coli* (1). To capture large, scarce, essential nutrients, the outer membrane displays high-affinity customized β-barrels for active transport of diverse ligands (2). The energy for the active transport across the essentially unenergized outer membrane comes from a complex of cytoplasmic membrane proteins, TonB, ExbB, and ExbD. This complex harvests the cytoplasmic membrane proton gradient and transforms it into mechanical energy which drives vectoral transport of ligands through TBDTs and into the periplasmic space (3) (Fig. 1). Because these transporters bind directly only to TonB and not ExbB or ExbD during the energy transduction, they have been termed TonB-dependent transporters (TBDTs) (4). There is some confusion in the historical literature whereby they were first called TonB-dependent receptors because they were initially identified as receptors for colicins and bacteriophages, now known to be opportunistic agents (5–7). They have also been called TonB-gated transporters, TGTs, (8) and ligand-gated porins, LGPs, (9). *Escherichia coli* K12 encodes nine different TBDTs mostly devoted to acquisition of iron by various means with one devoted to cobalamin transport. While *E. coli* has dedicated TBDTs for a variety of siderophores, enterochelin (a.k.a. enterobactin) is the single iron-chelating siderophore that *E. coli* synthesizes and excretes to capture iron from its environment [for a review see (10)]. The TBDT that provides for the recovery of iron-bearing enterochelin is FepA.

**Figure 1.**
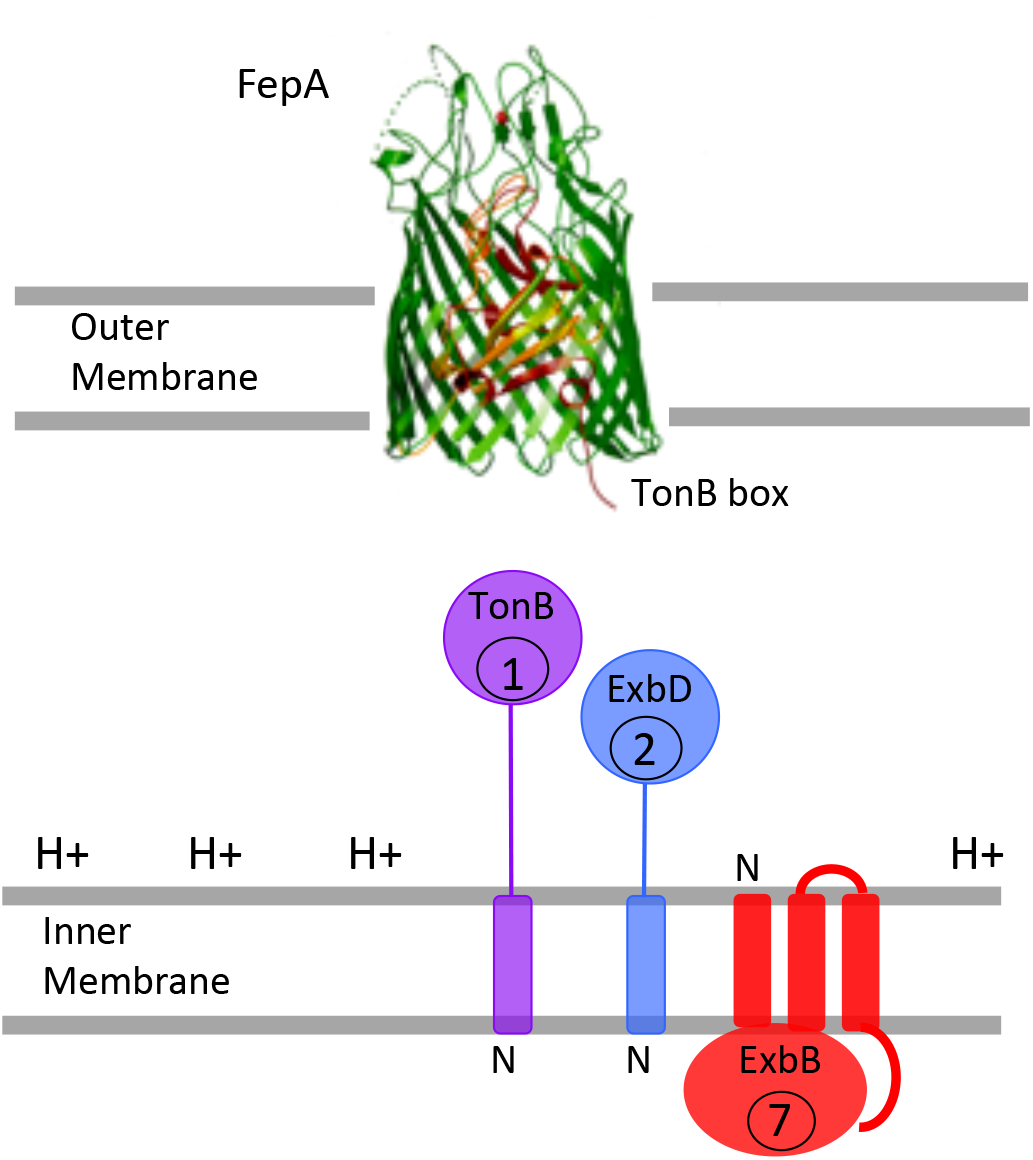
The TonB system of *Escherichia coli* K12. The TonB-dependent transporter, FepA, is shown in the outer membrane. At its extreme amino terminus, the TonB box, the only known site of *in vivo* interaction with TonB, is shown protruding into the periplasm. The topologies and cellular ratios of the cytoplasmic membrane proteins TonB, ExbB and ExbD are shown in the cytoplasmic membrane. The protonmotive force gradient of the cytoplasmic membrane (H+) is shown. The crystal structure of FepA was solved by Buchanan et al. (60).

Each TBDT consists of a 22-stranded β-barrel, the lumen of which is occluded by an essential internal globular domain of ∼ 150 residues called the cork (or hatch) [(11); for a review see (2)]. Because they are similar in structure, the results from one TBDT largely apply to most TBDTs. The mechanism by which TBDTs actively transport ligands such as the iron-siderophore enterochelin or cobalamin across the outer membrane remains a mystery, but there is general agreement that the cork must somehow move.

A TBDT cork has both a mechanically weak (approximately residues 1-70) and mechanically recalcitrant domain (approximately residues 70-150) both *in vivo* and *in vitro* (9, 12–14). An essential motif of five to seven mostly conserved residues known as the TonB box occupies the amino terminus of the mechanically weak domain. The TonB boxes of TBDTs are interchangeable, indicating that they do not mediate ligand specificity (15, 16).

The precise energy-transducing interaction of TonB with TBDTs has been challenging to define by structural determinations *in vitro* because ExbB and ExbD functions, and the protonmotive force of the cytoplasmic membrane are all required for TonB-dependent energy transduction. Furthermore, *in vitro* and *in vivo* TonB binds to TBDTs regardless of its ability to transduce energy, (17–22), suggesting that certain residue-specific interactions result in energy transduction while other interactions fail to accomplish it.

The TonB protein is anchored in the cytoplasmic membrane by its hydrophobic amino terminal signal anchor with the rest of the protein (residues 33-239 occupying the periplasmic space [(23), Fig.1]. Interestingly, there are no essential residues in TonB (17, 24–28). In fact, even the His20 residue in the transmembrane domain can be replaced with non-protonatable Asn and retain full function, suggesting that the H20A mutation used in that study renders TonB inactive through steric distortion of its transmembrane domain in complex with ExbB and ExbD transmembrane domains (28).

There are, however, seven residues in the periplasmic TonB carboxy terminus (out of 90 sequentially scanned) that are functionally important (Y163, F180, G186, F202, W213, Y215, and F230) (27). With the exception of G186, these residues represent the complete set of aromatic residues in the last 90 residues of the carboxy terminus from 150-239, with the only other aromatic residue in the entire periplasmic domain (residues 33-239) being F125. We think these seven residues are the key because:

1) When substituted with Ala or Cys, each of these residues exhibits an idiosyncratic phenotype, with the profile of activities in four different assays being distinct for each of the seven substitutions (27, 29). They appear to be the means by which TonB discriminates among different TBDTs or possibly the colicins that parasitize them (30). In contrast, Cys substitution at the only other aromatic residue, TonB F125C, supports wild-type activity.

2) They are synergistic with one another such that any combination of two mutations (2 Ala, 2 Cys, or a combination) is completely inactive in all assays in a double mutant cycle analysis. For example, TonB F202A, W213A used in previous studies is completely inactive, whereas TonB F125A is not synergistic with substitutions at any of the seven residues (17, 27, 29). It therefore seems to set a maximal boundary on the active domain of the TonB carboxy terminus from G186 to F230, which contains a single amphipathic helix (residues 199-216).

3) Cys substitutions in five out of the seven important carboxy terminal residues (G186C, F202C, W213C, Y215C, F230C) are the only ones out of the 90 Cys substitutions that form disulfide-linked triplet homodimers (17, 27). While both inactive and active TonB binds to transporters, the disulfide-linked homodimers formed through the five Cys substitutions are trapped in configurations such that they no longer fractionate significantly with the outer membrane (17, 27). In contrast, TonB F125C, which appears to be outside the active domain of the carboxy terminus, forms triplet homodimers that, like wild-type TonB, still fractionate ∼ 40% with the outer membrane, indicating that the subsequent, more carboxy-terminal residues, including especially the amphipathic helix, are free to undergo necessary conformational changes (31).

TonB is the limiting protein in the TonB system (32) and different TBDTs must compete for its attention (33). TonB therefore interacts transiently with ligand-loaded TBDTs in *E. coli* K12, giving rise to an energy transduction cycle (67). Over the years we have defined stages in that cycle *in vivo*, the model for which is depicted in Fig. 2.

**Figure 2.**
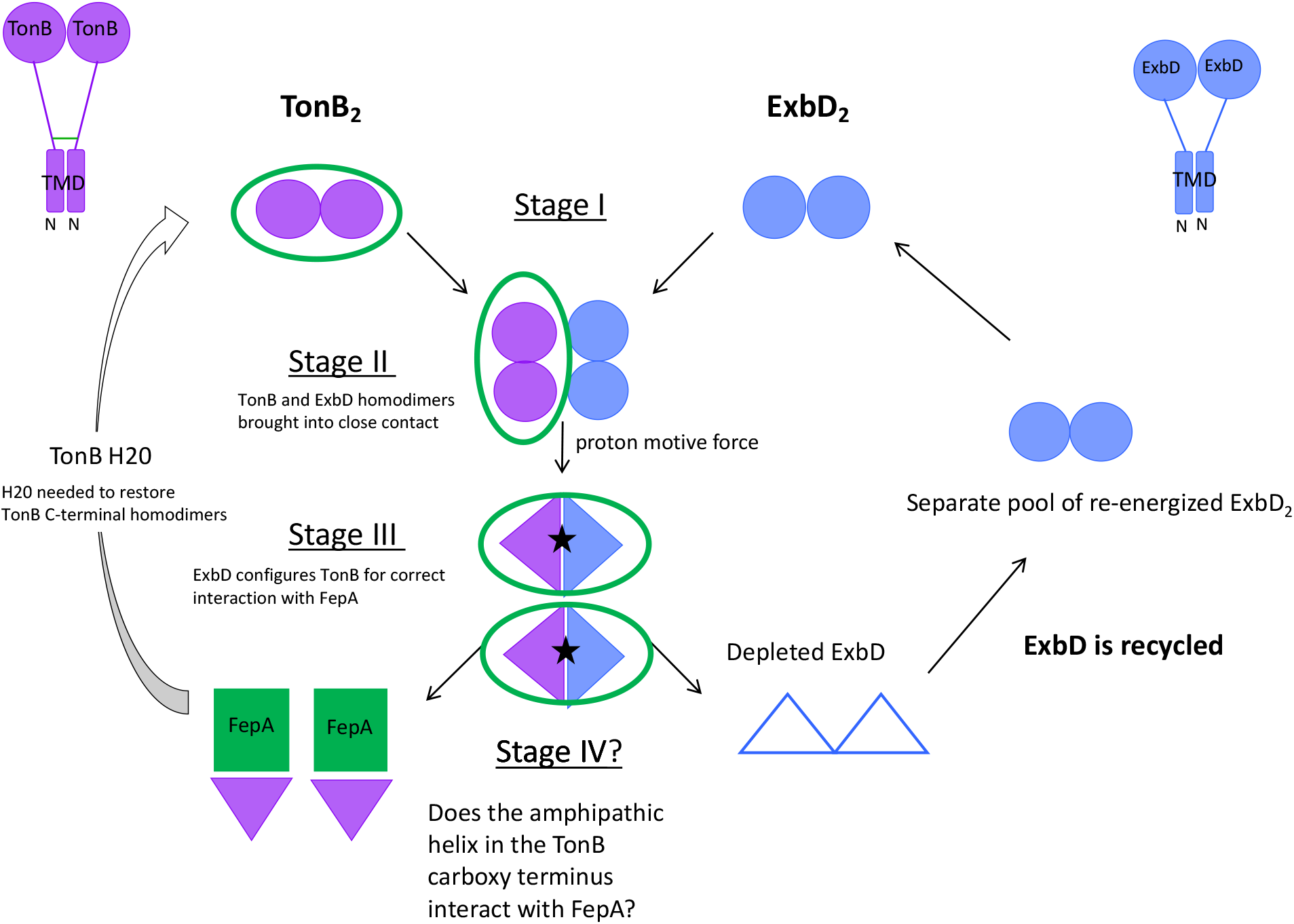
Key *in vivo* interactions of the TonB carboxy terminus during the energy transduction cycle involve the amphipathic helix (residues 199-216) [adapted from (31).]. Full length TonB (purple, upper left corner) and ExbD (blue, upper right corner) each have a single transmembrane domain signal anchor in the cytoplasmic membrane with the bulk of the residues occupying the periplasmic space. Filled purple circles/triangles are TonB carboxy termini. Filled blue circles/triangles are ExbD carboxy termini. The TonB amino terminal domain remains homodimerized throughout the cycle, with its periplasmic carboxy terminus undergoing sequential protein-protein interactions (31). Interactions of the periplasmic carboxy-terminal domains of both TonB_2_ and ExbD_2_ homodimers are shown. In Stage I, H20 in the TonB transmembrane domain is required for TonB carboxy termini to form obligatory homodimers through residues in and near the carboxy terminal amphipathic helix (residues 199-216), (17, 31). ExbB tetramers (ExbB_4_) independently stabilize both TonB_2_ and ExbD_2_, homodimerized through their carboxy termini, and are proposed to be the scaffolds upon which TonB_2_ and ExbD_2_ are independently assembled (ExbB_4_ is not shown). In Stage II, TonB_2_ and ExbD_2_ homodimers are brought into close contact by ExbB tetramers but have not yet formed the heterodimers of the subsequent Stage. In Stage III, in the presence of the cytoplasmic membrane protonmotive force (PMF), the TonB_2_ and ExbD_2_ carboxy termini reassort to form two TonB-ExbD heterodimers. ExbD, which contains the sole potentially PMF-responsive residue (Asp25) among five transmembrane domains that make up the TonB system, configures TonB correctly for a productive interaction with FepA [(28, 74); Jana and Postle, unpublished]. This is necessary because inactive TonB also binds to TBDTs but without energy transduction, meaning that the correct configuration must be based on prior interaction with ExbD. Stage IV is binding of a monomeric carboxy terminus of TonB to FepA, such that active transport of the siderophore enterochelin across the outer membrane occurs. Notably, the TonB amphipathic helix makes important contacts with another TonB or ExbD in Stages I-III (green circles), however it has never been tested for important contacts with FepA (green square). After a transport event, H20 is required for re-formation of TonB dimers in Stage I. ExbD_2_ is de-energized after this event (empty blue triangles) and needs to be recycled. We speculate that ExbD_2_ moves in and out of the complex escorted by ExbB tetramers. A separate pool of recycled ExbB_4_-ExbD_2_ is hypothesized to exist to replenish Stage I ExbB_4_-ExbD homodimers. See (31) for a full explanation of the experimental basis for the model.

In the model, the TonB carboxy termini of homodimers are conformationally dynamic while the amino termini remain stably homodimerized throughout the energy transduction cycle (31). Protonmotive force of the cytoplasmic membrane is transduced into active transport at the outer membrane through sequential contacts by the TonB carboxy terminus, first with itself, then with the ExbD carboxy terminus, then with a TBDT (31). *In vivo* interaction sites between TonB homodimers and between TonB-ExbD heterodimers have been identified, and a common region between them is the TonB amphipathic helix (residues 199-216; Fig. 3) (27, 34, 35). The primary goal of this study was to determine if the TonB amphipathic helix played a role in contact with FepA.

**Figure 3.**
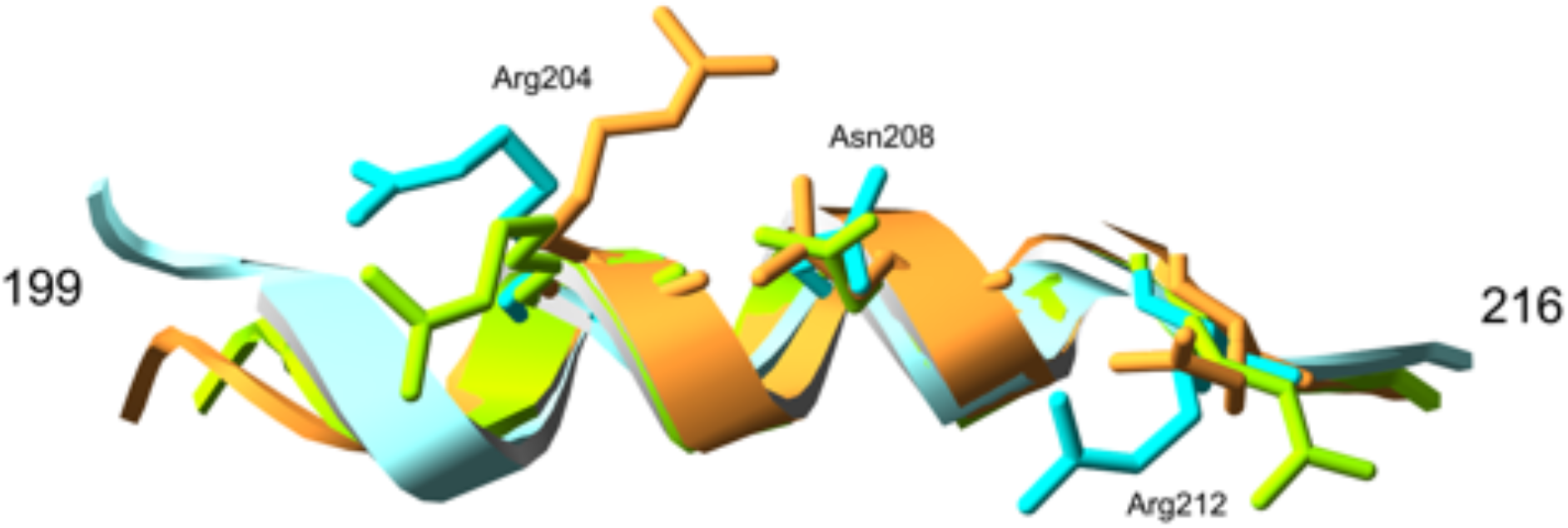
Superimposition of the TonB amphipathic helices from TonB crystal structures. Blue is from Chang et al. (75); green is from Peacock et al. (76); gold is from Shultis et al. (45). The hydrophilic residues that, as Cys substitutions, interact with the FepA TonB box Cys substitutions in Fig. 7 are shown.

Here we explored TonB interactions with three different regions of FepA— the TonB box, the cork, and the β-strand turns of the barrel. A novel *in vivo* interaction between Cys substitutions in the hydrophilic face of the essential TonB amphipathic helix and Cys substitutions in the essential FepA TonB box was identified. Interactions between TonB amphipathic helix and β-strand turns of the FepA barrel were identified, providing the first *in vivo* support for a surveillance model where TonB searches for a TBDT. The difference in interaction profiles with the mechanically weak domain of the FepA cork between the TonB amphipathic helix and TonB Q160 led to a model where the amphipathic helix pulls the mechanically weak domain of the FepA cork out of the barrel sufficiently that TonB Q160 interacts with otherwise buried residues.

## RESULTS

### The carboxy terminal TonB amphipathic helix is essential for TonB system activity

The TonB region from ∼ R158-N162, centered on TonB Q160 interacts with the TonB boxes of TBDTs *in vivo* [Fig. 4A; (4, 36)]. While the TonB box as a whole is essential for TBDT activity, its precise amino acid composition is tolerant of substitutions except for structure-breaking residues such as L8P in BtuB or I14P in FepA, substitutions that result in inactivation of the TBDT (4, 15, 36, 37). Even then, the TonB Q160 region still interacts with the BtuB L8P mutant TonB box *in vivo*, but in a different way (4).

**Figure 4.**
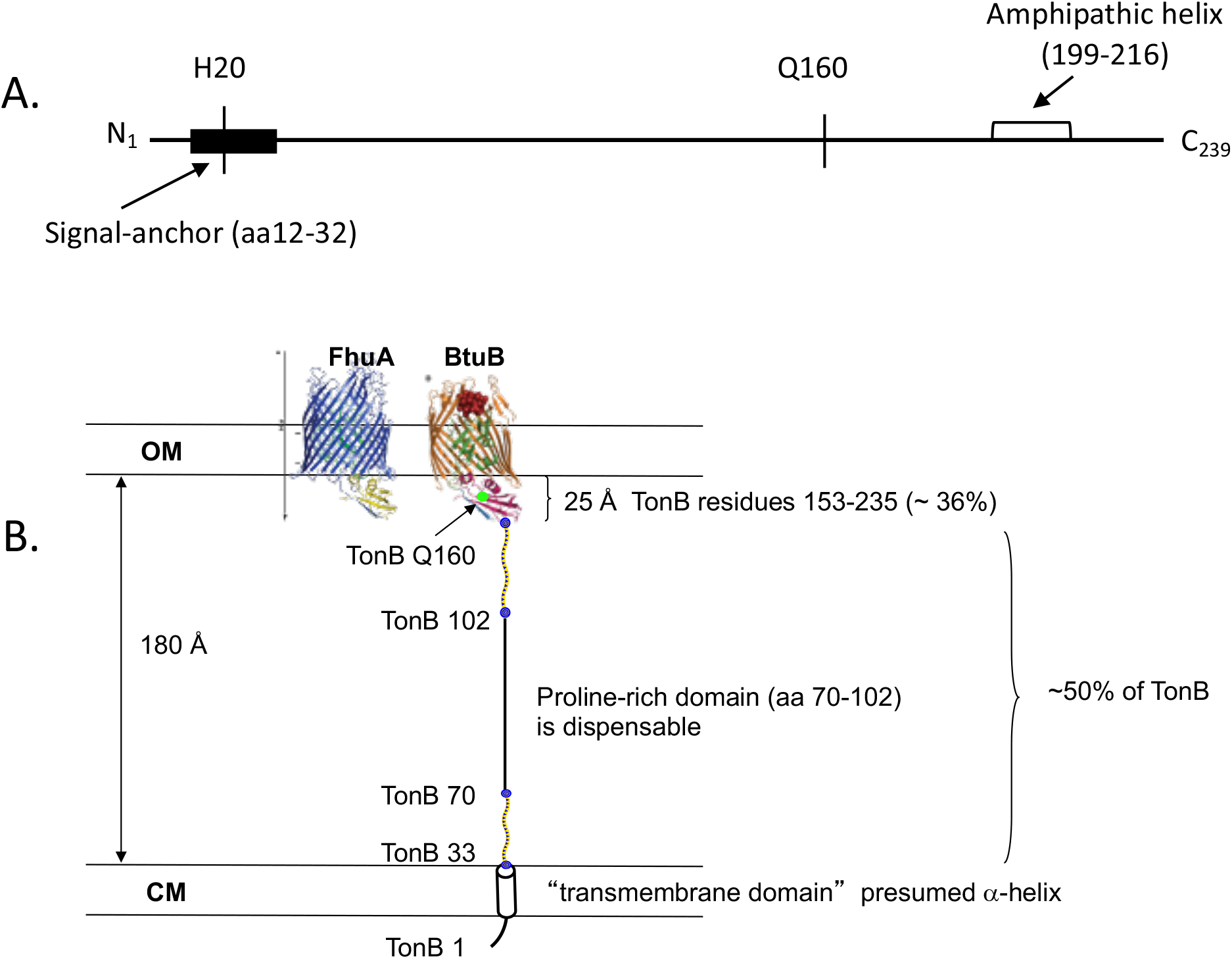
TonB protein information. **(A)** Relevant features of the TonB primary amino acid sequence (residues 1-239) are shown. H20 in the TonB transmembrane signal anchor domain renders TonB inactive when substituted with alanine (44). Locations of TonB Q160 and the amphipathic helix are shown. (**B)** The conformation of TonB predicted by the crystal structures of its carboxy terminus would be unable to reach the TBDTs. TonB residues 33–69 and 103–149, predicted to be intrinsically disordered regions (44) are depicted as yellow rectangles. TonB Q160 is the green dot within the structured carboxy terminus of the BtuB-TonB structure. The proline-rich domain, which contributes ∼ 100 Å to the extension of TonB across the periplasmic space (38), can be deleted without inactivating TonB (24, 39). The span of the periplasmic space was estimated based on crystal structure reconstructions of the AcrA/B/TolC complex which has proteins in both outer and cytoplasmic membranes (77). The crystal structure of FhuA-TonB is from Pawelek et al. (20). The crystal structure of BtuB-TonB is from Shultis et al. (45). Part B of this figure is from (44). Abbreviations: CM, cytoplasmic membrane; OM, outer membrane.

Based on the position of residue Q160 within the TonB carboxy terminus, the solved crystal structures of TBDTs, and the solved crystal structures of TonB carboxy termini in complex with those TBDTs (Fig. 4B), it seems unlikely that Q160 contacts a TBDT without significant conformational changes to make the TonB box more accessible. The deletion of the proline rich domain that accounts for ∼ 100 Å of TonB’s reach to a TBDT has negligible effect upon its activity unless the periplasmic space is artificially expanded by transient exposure to high salt (24, 38, 39). This observation suggests that residues nearer the carboxy terminus of TonB could be more important. There is also evidence that unknown regions in addition to the FepA TonB box and TonB Q160 are involved in transport (12, 40). Furthermore, the TonB Q160 region is not essential, suggesting that its role in contacting a TBDT is not essential (11).

The periplasmic TonB carboxy terminus contains an amphipathic helix (residues 199-216) which has been intriguing for many years (24, 29, 41). It includes three of the seven functional carboxy terminal residues, F202, W213 and Y215. To further explore the role the amphipathic helix plays in the mechanism of TonB-dependent energy transduction, we deleted the amphipathic helix codons 199-216 from plasmid pKP325, resulting in plasmid-encoded TonB_ΔAH_ (Fig. 5A). Plasmids expressing chromosomal levels of TonB_ΔAH_ (pKP476) were unable to complement KP1477 (Δ*tonB*) in cross-streaks against colicins B, Ia, and M, the most sensitive assays known for TonB function, requiring ∼1 active TonB molecule per cell (42).

**Figure 5.**
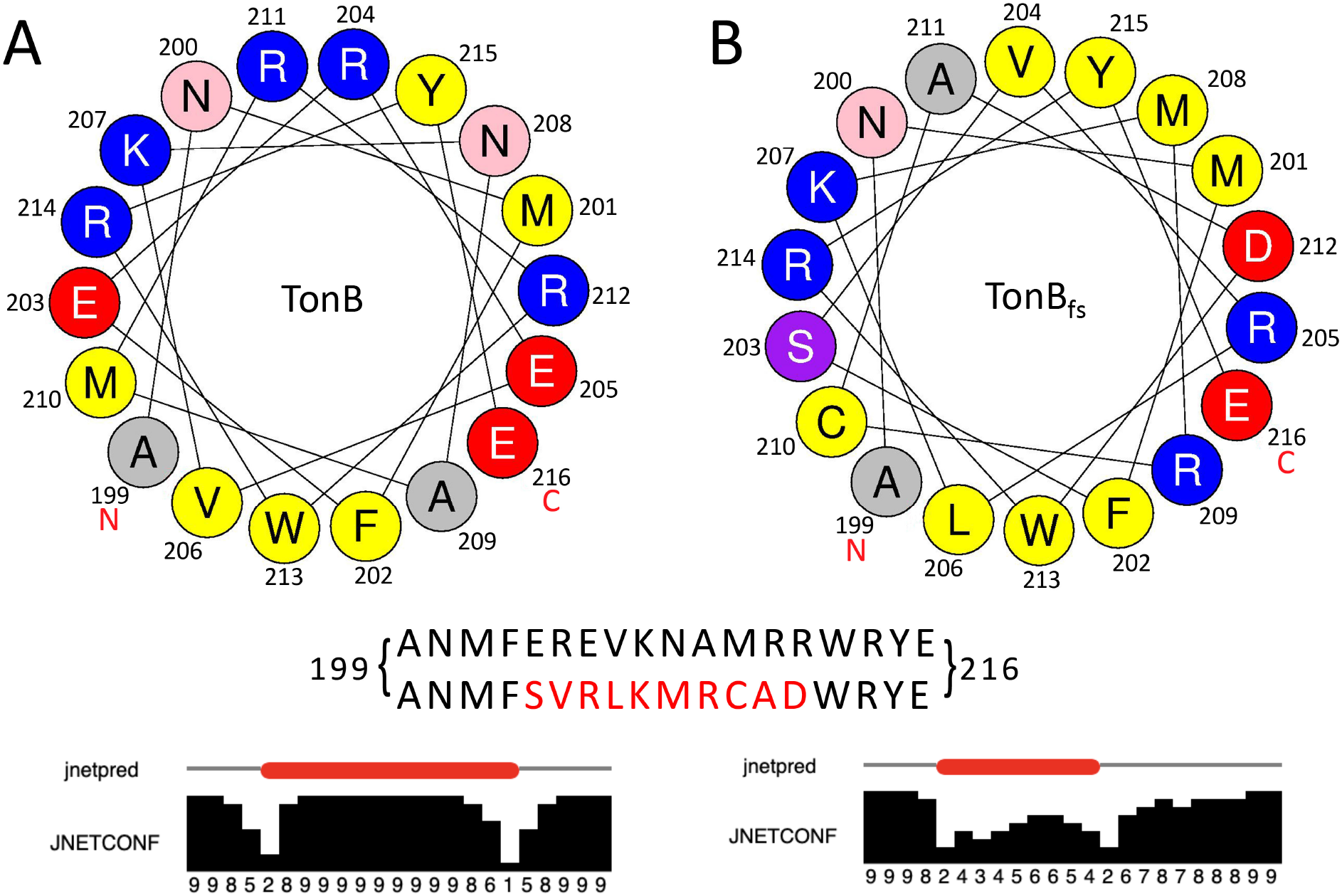
Comparison of wild type TonB and frame-shifted (TonB_fs_) amphipathic helix regions (residues 199-216). **(A)** Helical wheel diagram of the TonB amphipathic helix and its corresponding JPRED4 prediction (bottom) as compared to (**B)** the corresponding frame-shifted region of TonB presented in a helical wheel diagram with its corresponding JPRED4 prediction (bottom). For JNetPred, predicted helices are shown in red. For JNETCONF, high values along the bottom edge indicate high confidence in the prediction (78). The comparison of the two amino acid sequences is shown in the middle panel, with the frame-shifted residues noted in red. The frameshifted version has lost much but not all of its helical character and it resulted in multiple substitutions in the core helix residues 203-212.

TonB_ΔAH_ fractionated on sucrose density gradients with ∼ 60% located in the cytoplasmic membrane fractions and ∼40% with the outer membrane fractions (Fig. 6A), the same proportions with which wild-type TonB fractionates (43). It is not known what causes TonB to bind sufficiently tightly to outer membrane components that ∼ 40 % is pulled out of its complex with cytoplasmic membrane proteins ExbB and ExbD to fractionate with the outer membrane. One hypothesis is that TonB outer membrane fractionation reflects a transient tight association with outer membrane molecules--likely TBDTs--during Stage IV in the energy transduction cycle (Fig. 2). While the region required for outer membrane fractionation, residues 175-239, includes the amphipathic helix (43) and is responsible for the ability of TonB to formaldehyde crosslink to FepA (37), this result indicated that the amphipathic helix was not the region responsible for fractionation of TonB with the outer membrane.

**Figure 6.**
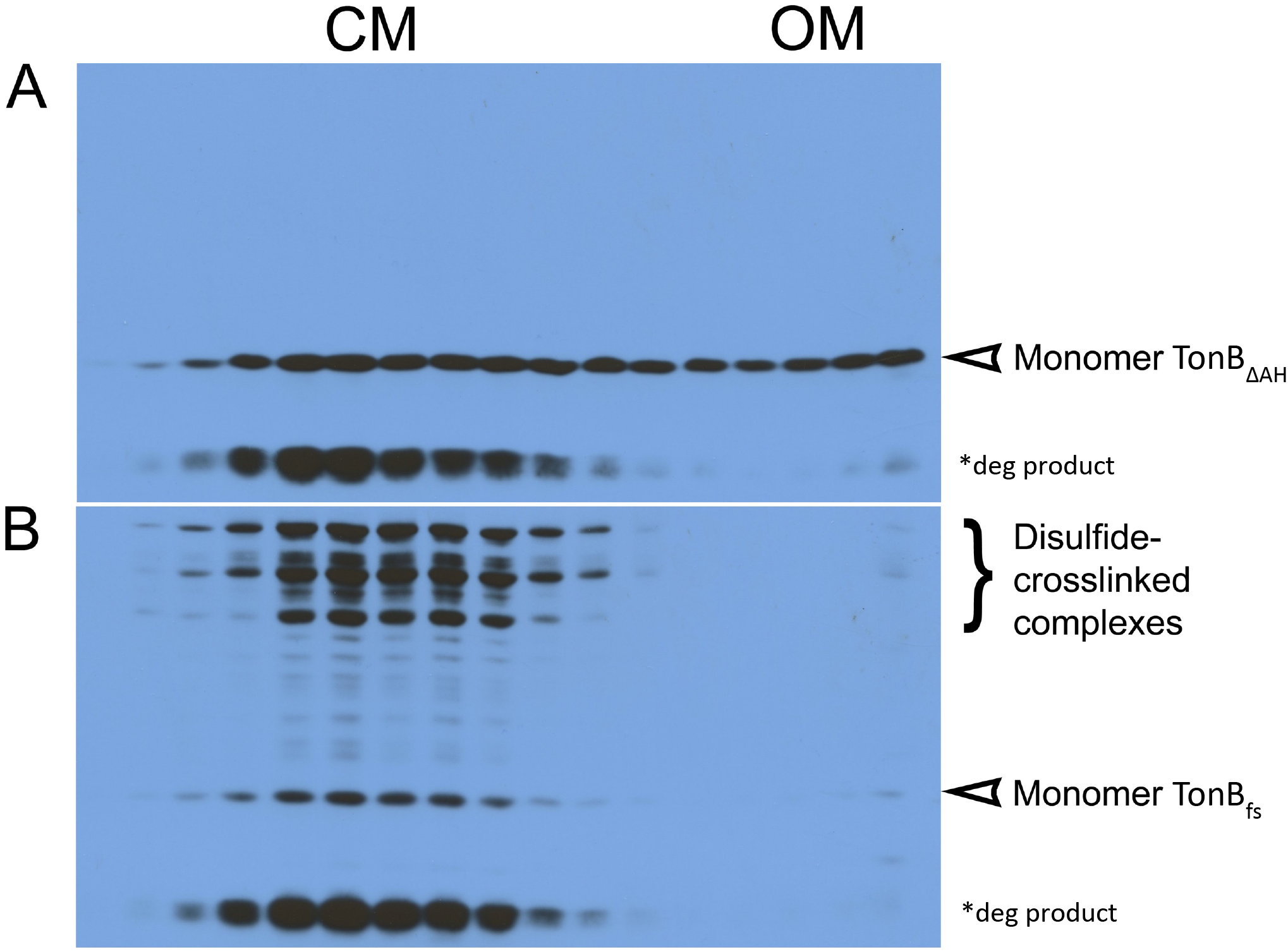
Monomeric TonB with major alterations of its amphipathic helix retains its ability to associate with the outer membrane. Wild-type TonB fractionates ∼ 40% with the outer membrane and ∼ 60% with the cytoplasmic membrane (43). (**A)** Sucrose density gradient fractionation of TonB_ΔAH_, where TonB associates with both membranes. (**B)** Sucrose density gradient fractionation of TonB_fs_. Because most of the TonB_fs_ is trapped as triplet homodimers through M210C that cannot associate with the outer membrane (17, 31), only a small amount of monomer is apparent in OM fractions on this exposure. On longer exposures, monomer TonB_fs_ association with the outer membrane becomes more apparent (data not shown). CM is for cytoplasmic membrane fractions; OM is for outer membrane fractions. Immunoblots of SDS polyacrylamide gels with anti-TonB monoclonal antibody are shown (71).

The inactivity of TonB_ΔAH_ could have been due to structural perturbation. Alternatively, it could have been due to the simultaneous deletion of residues F202, W213, and Y215, since combination of two Ala substitutions at any of those residues renders TonB inactive (27, 29). Individual Cys substitutions from E203 through R212 have little phenotypic effect (27). To retain F202, W213, and Y215 and broadly restore the overall structure, we shifted the 10-amino acid core region within the helix out of frame starting at residue 203 and shifted it back into frame at residue 213 (TonB_fs_, pKP372). The result for TonB_fs_ was that the predicted helical region was shortened slightly to encompass residues 201-209 (as analyzed by JPRED), lost much of its amphipathic character, and at residue 210, Met was substituted with Cys due to the frameshift (Fig. 5). Like TonB_ΔAH_, TonB_fs_ was completely insensitive in the colicin assays.

Due to the newly created M210C, TonB_fs_ efficiently formed triplet homodimers. Like those of the Cys substitutions in functionally important residues, the TonB_fs_ triplet homodimers fractionated essentially entirely with the cytoplasmic membrane, suggesting that they had trapped TonB at a stage in the energy transduction cycle before TonB associates with the outer membrane [Fig. 6B; (17, 27)]. TonB M210C in the context of otherwise wild-type residues is active and does not form triplet homodimers (27).

The *in vivo* dynamics of the TonB carboxy terminus suggest that it achieves a monomeric conformation at some point in the energy transduction cycle to allow productive interaction with FepA (31). Because a detectable proportion of TonB_fs_ remained monomeric and was also found in the outer membrane on longer exposures (data not shown), its inactivity was not due to 100% entrapment as a triplet homodimer (Fig. 5B). We concluded that one or more aspects of the core amphipathic helix domain were required for TonB activity.

### The TonB amphipathic helix interacts functionally with the FepA TonB box

An analysis of whether and how the TonB amphipathic helix interacts *in vivo* with any of the TBDTs has never been performed. To investigate interactions between the TonB amphipathic helix and the FepA TonB box, we used *in vivo* disulfide crosslinking of TonB and FepA Cys substitutions expressed at chromosomal levels, followed by electrophoresis on non-reducing SDS gels and immunoblotting with anti-TonB monoclonal antibodies.

We surveyed crosslinking by TonB amphipathic helix residues R204C, V206C, N208C, A209C, and R212C (Fig. 5). When expressed at chromosomal levels, each of the TonB Cys substitutions was at least 60% active in ^55^Fe-ferrichrome transport assays (Table 1). TonB Q160C, over 100% active and a known site of interaction with other TBDT TonB boxes, was also tested for comparison to the amphipathic helix Cys substitutions [Table 1; (4, 16)]. The FepA TonB box Cys substitutions tested were D12C, T13C, I14C, V15C, V16C and T17C; FepA T13C is fully active (44) as was FepA V16C (Table 1) with the rest assumed to be active.

**Table 1:**
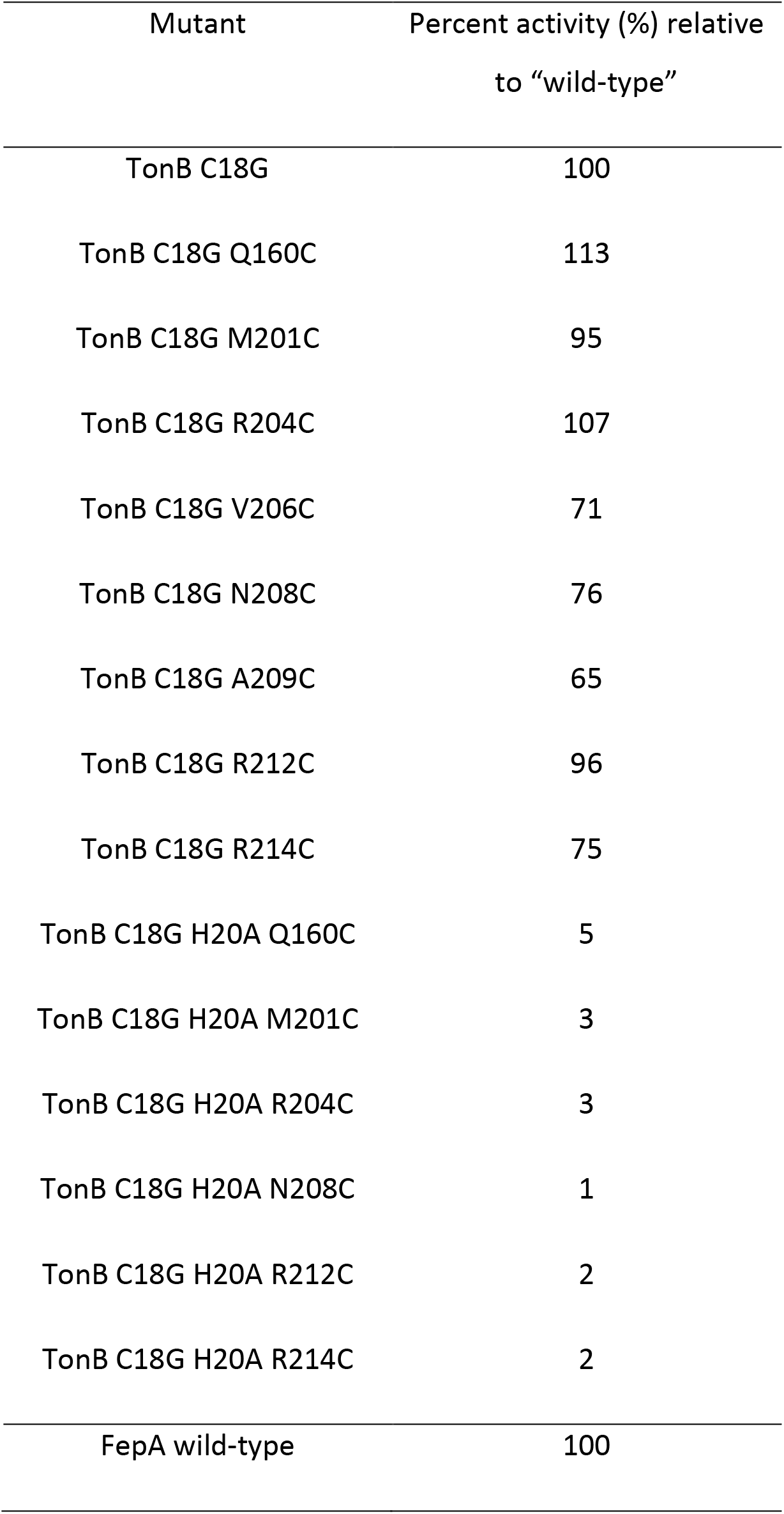
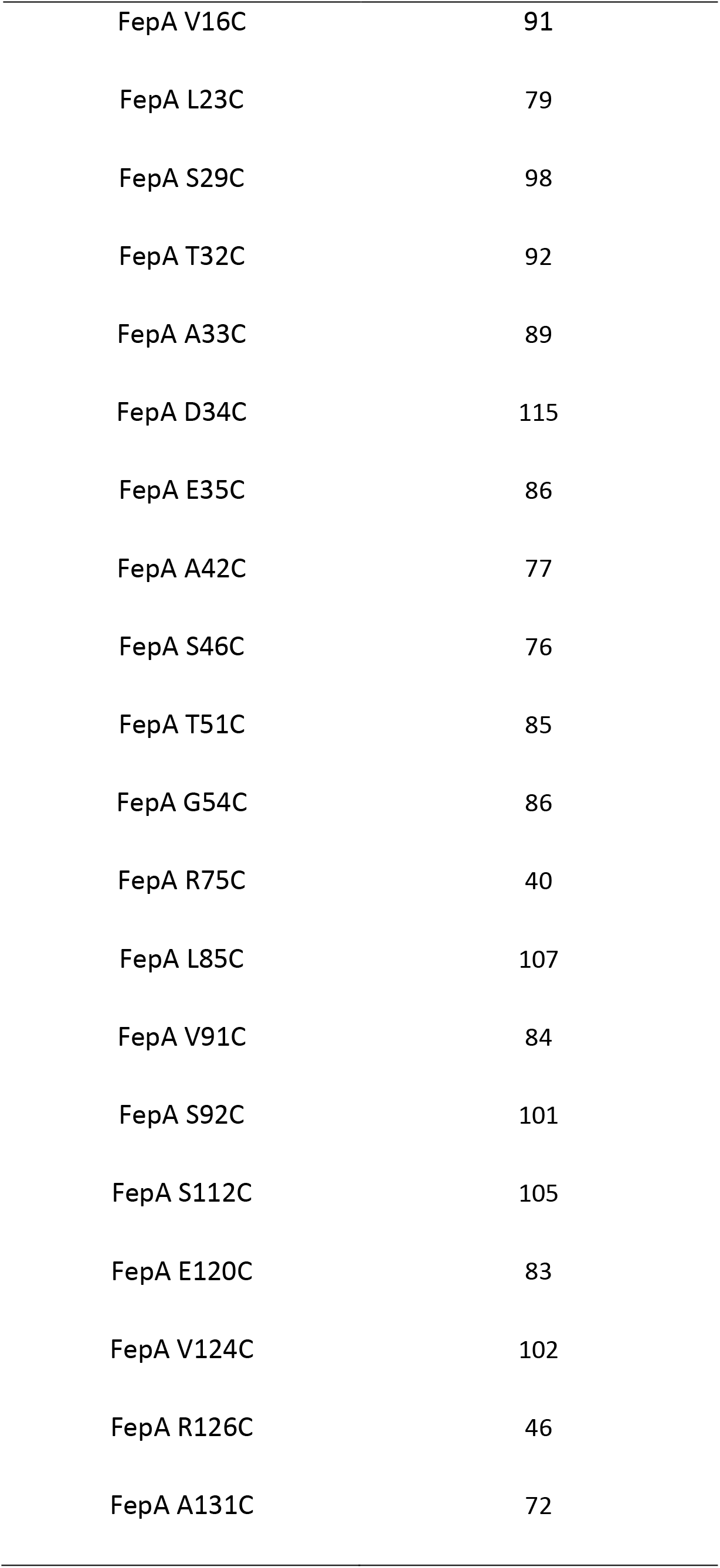
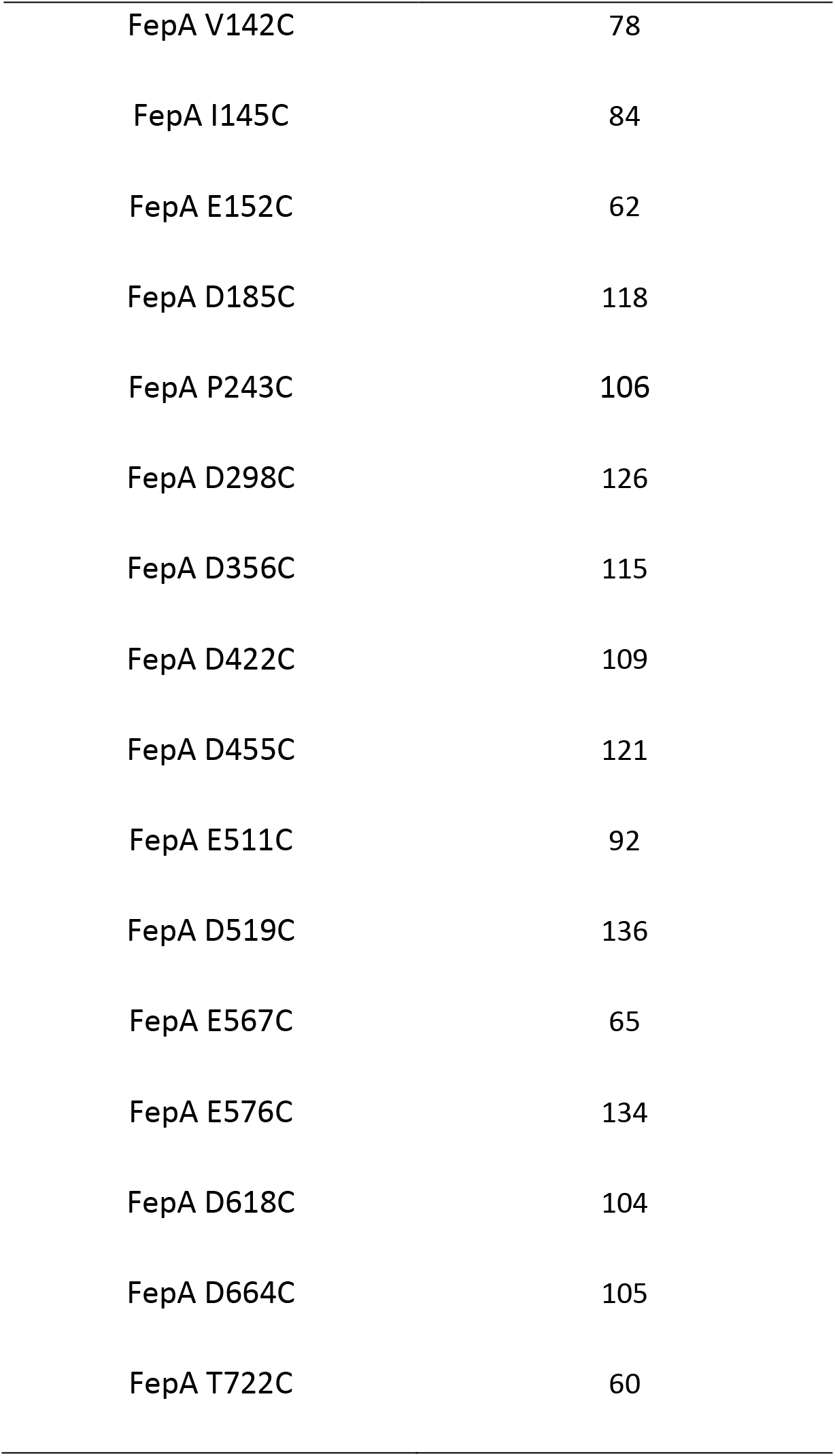
[^55^Fe]-enterochelin transport activities of TonB and FepA Cys substitutions.

| Mutant | Percent activity (%) relative<br>to “wild-type” |
| --- | --- |
| TonB C18G | 100 |
| TonB C18G Q160C | 113 |
| TonB C18G M201C | 95 |
| TonB C18G R204C | 107 |
| TonB C18G V206C | 71 |
| TonB C18G N208C | 76 |
| TonB C18G A209C | 65 |
| TonB C18G R212C | 96 |
| TonB C18G R214C | 75 |
| TonB C18G H20A Q160C | 5 |
| TonB C18G H20A M201C | 3 |
| TonB C18G H20A R204C | 3 |
| TonB C18G H20A N208C | 1 |
| TonB C18G H20A R212C | 2 |
| TonB C18G H20A R214C | 2 |
| FepA wild-type | 100 |
| FepA V16C | 91 |
| FepA L23C | 79 |
| FepA S29C | 98 |
| FepA T32C | 92 |
| FepA A33C | 89 |
| FepA D34C | 115 |
| FepA E35C | 86 |
| FepA A42C | 77 |
| FepA S46C | 76 |
| FepA T51C | 85 |
| FepA G54C | 86 |
| FepA R75C | 40 |
| FepA L85C | 107 |
| FepA V91C | 84 |
| FepA S92C | 101 |
| FepA S112C | 105 |
| FepA E120C | 83 |
| FepA V124C | 102 |
| FepA R126C | 46 |
| FepA A131C | 72 |
| FepA V142C | 78 |
| FepA I145C | 84 |
| FepA E152C | 62 |
| FepA D185C | 118 |
| FepA P243C | 106 |
| FepA D298C | 126 |
| FepA D356C | 115 |
| FepA D422C | 109 |
| FepA D455C | 121 |
| FepA E511C | 92 |
| FepA D519C | 136 |
| FepA E567C | 65 |
| FepA E576C | 134 |
| FepA D618C | 104 |
| FepA D664C | 105 |
| FepA T722C | 60 |

The amphipathic helix substitutions at R204C, N208C and R212C formed disulfide-linked complexes with all FepA TonB box Cys residues except N208C with FepA T17C (for unknown reasons) (Fig. 7A). In contrast, TonB V206C and A209C did not form significant complexes with any of the FepA TonB box Cys residues, thus constituting a non-reactive hydrophobic face of the amphipathic helix. Steady state levels of monomeric TonB from samples in Fig. 7A are presented in Fig. 7B.

**Fig. 7.**
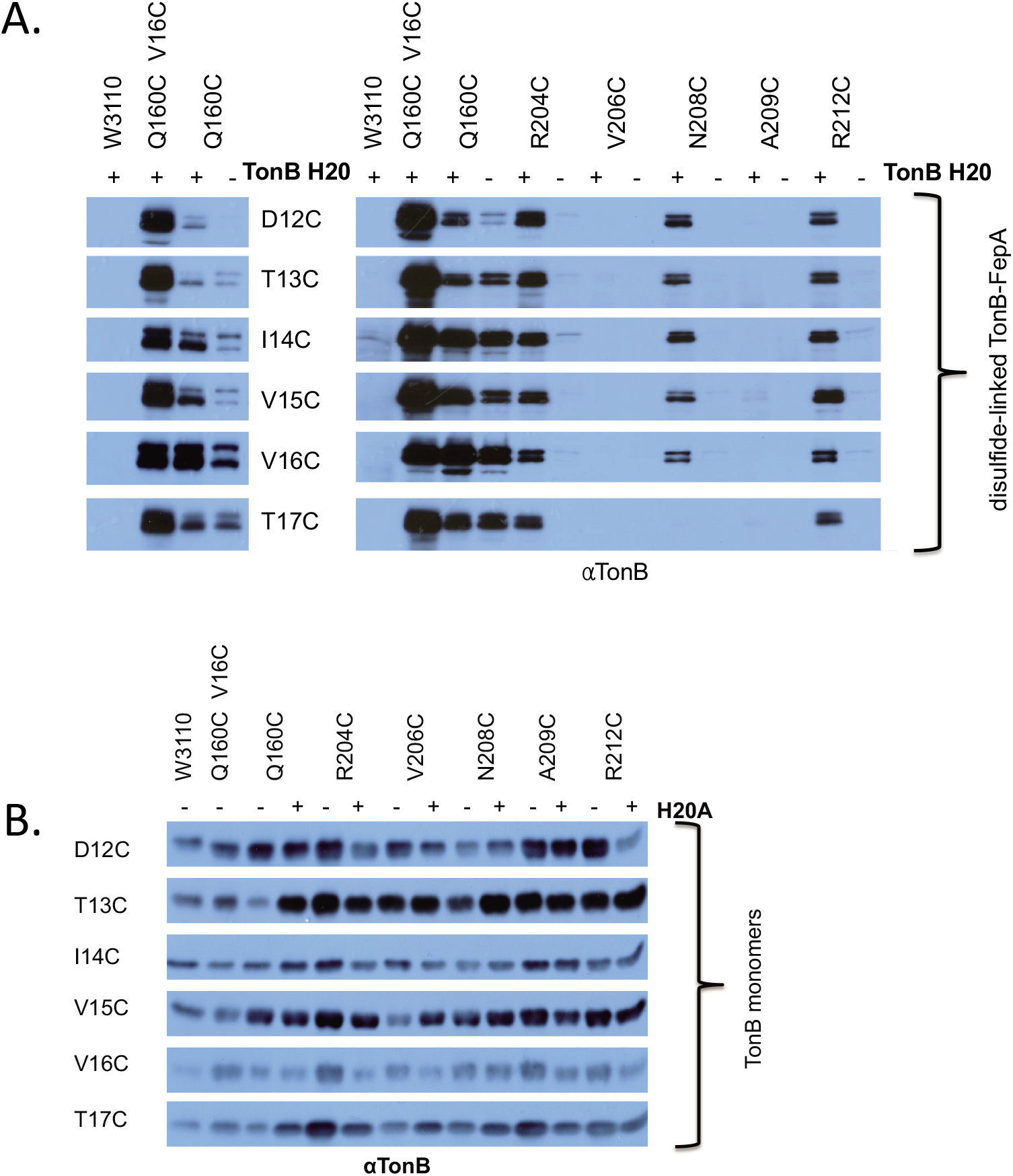
Cys substitutions in the essential TonB amphipathic helix form *in vivo* disulfide crosslinks with FepA TonB box Cys substitutions. Strain KP1491 [W3110 Δ*fepA,* Δ(*tonB,P14*)*::kan*] with various TonB and FepA plasmid combinations was grown and analyzed as described in Materials and Methods. TonB and its Cys substitutions are indicated across the top of the immunoblots. FepA Cys substitutions in the TonB box are indicated between the two panels (in (A) or on the left side of the immunoblot in (B). (+) indicates the presence of the wild-type H20 allele in the TonB transmembrane domain. (-) indicates the presence of the inactivating H20A mutation. Immunoblots of the ∼ 116 kDa region of non-reducing SDS polyacrylamide gels developed with monoclonal anti-TonB antibody are shown. **(A)** Wild-type control W3110 shows that wildtype TonB and wildtype FepA do not innately form stable complexes. Panel right: TonB-FepA disulfide-linked complexes are shown. Panel left: shorter exposures of the first four lanes of panel right are shown. In panel left, the TonB Q160C-FepA V16C pair demonstrated the most efficient crosslinking among five TonB FepA TonB box Cys substitutions tested and was therefore used as a standard for relative levels in all subsequent figures characterizing disulfide crosslinks. **(B)** Steady state levels of chromosomally encoded TonB in W3110 and plasmid-encoded TonB variants from samples in (A) are shown as immunoblots of the ∼ 36 kDa region of reducing SDS polyacrylamide gels developed with anti-TonB monoclonal antibody.

To determine if the TonB-FepA disulfide crosslinks were biologically relevant, each TonB Cys substitution was also paired with the H20A mutation in the TonB transmembrane domain (44). This mutation inhibits homodimerization of TonB through its carboxy terminus, a step necessary for formation of carboxy terminal TonB-ExbD heterodimers and the subsequent correct interaction of the TonB carboxy terminus with FepA [Figs. 2 and 4A, (18, 31)]. TonB H20A epitomizes the behavior of inactive TonB because it still interacts at unknown sites with FepA *in vivo* (17, 18). The TonB H20A mutation rendered all the TonB Cys substitutions inactive (Table 1).

The H20A mutation essentially eliminated complex formation by R204C, N208C and R212C, indicating that the H20 wild-type versions were engaging in biologically relevant interactions. The possibility that H20A somehow promoted new interactions with the hydrophobic face (substitutions V206C and A209C) was excluded since no complexes were observed. These results indicated that the TonB amphipathic helix contacted the essential FepA TonB box *in vivo*, consistent with its role in TBDT reception of TonB-transmitted energy. The TonB amphipathic helix is the first known alternative to the TonB Q160 region for contact with the FepA TonB box.

In the solved co-crystal structures of the TonB carboxy terminus with the TBDTs BtuB and FhuA, TonB residues R204, N208 and R212 of the amphipathic helix interact with the barrels, but not the TonB boxes [Fig. 8, (20, 45)]. The lack of interaction with the TonB amphipathic helix in those elegant co-crystal structures supports the idea that the TonB carboxy terminus remains able to bind to TBDTs and other proteins in an “un-energized” conformation (17, 18, 46, 47).

**Figure 8:**
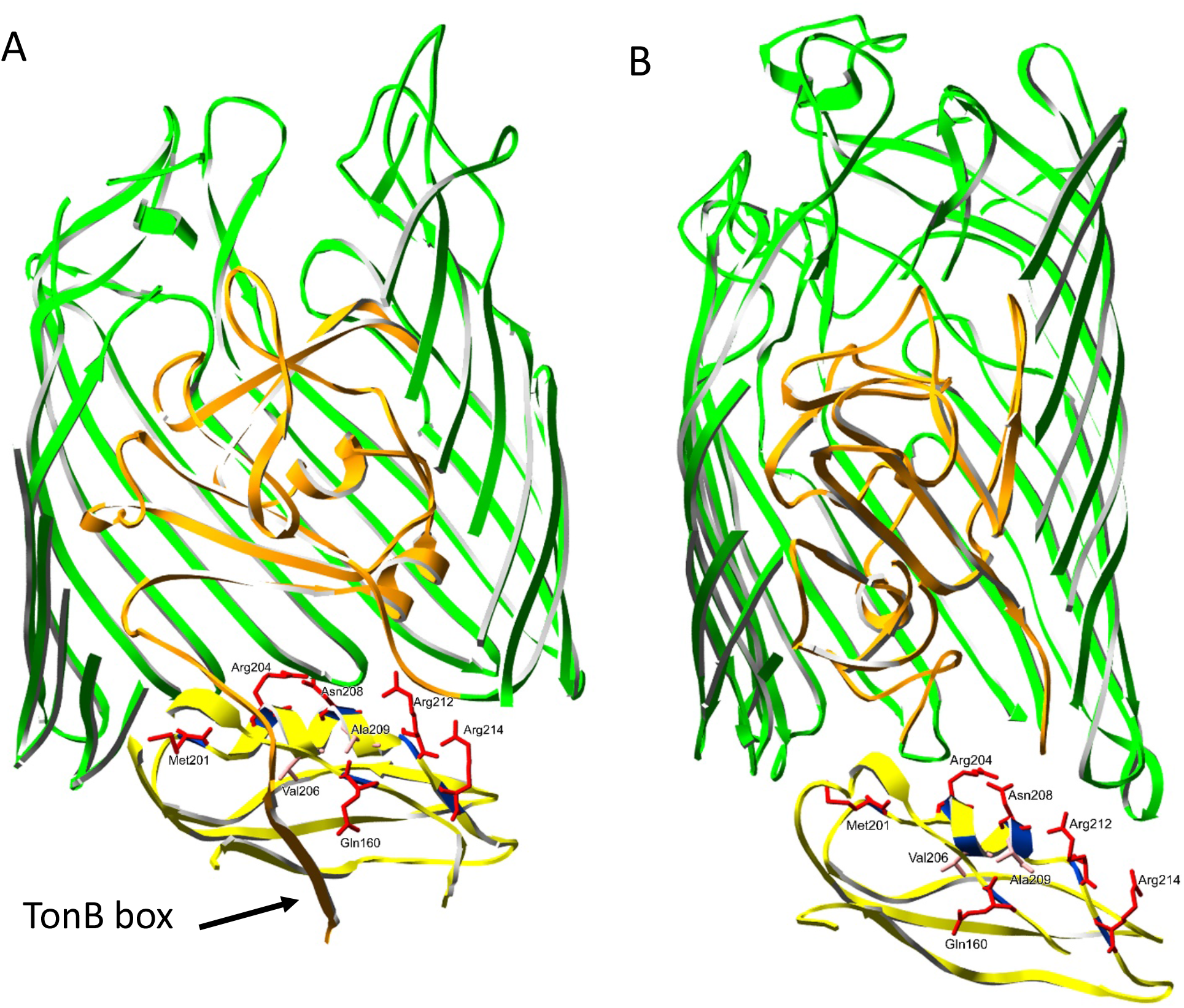
Ribbon diagrams of TonB Cys-substituted residues displayed on TonB carboxy terminus co-crystal structures with BtuB. [**A**; Shultis et al., (45)] **and FhuA** [**B**; Pawelek et al., (20)]. TonB Cys substitutions from this study that made disulfide crosslinks with a variety of Cys substitutions in FepA, including the TonB box, are in red; those in pink (A206C and A209C) made no crosslinks. The TonB box of BtuB is shown. The FhuA TonB box was not visible in the FhuA-TonB structure. The authors of that study propose that FhuA TonB box residues I9, T10, V11, and A13 interact with TonB residues V225, V226, L229, and K231 on one side and with TonB Q160 on the other (20).

### TonB Q160 interaction with the FepA TonB box is only partially dependent on TonB activity

Although the interaction of the TonB Q160 region with the TonB boxes of several TBDTs has been well-documented both *in vivo* and *in vitro* (4, 15, 16, 48–51), there has not yet been an analysis of how TonB Q160 interacts with the TonB box of FepA nor an analysis of effects of inactive TonB upon any TBDT TonB box interaction. In Fig. 7A, TonB Q160C crosslinked with all of the FepA TonB box Cys substitutions, consistent with its behavior seen previously for BtuB Cys substitutions in the TonB box (4). FepA V16C was chosen as the standard in all subsequent experiments because it exhibited the highest degree of disulfide crosslinking to TonB Q160C (Fig. 7A, left panel, lane 3), allowing comparisons of relative levels of TonB-FepA complex formation.

The H20A mutation detectably decreased TonB Q160C crosslinking, but without obliterating it altogether, suggesting that it represented a partially functional interaction (Fig. 7A, left panel, compare lanes 3 and 4).

All of the interactions gave rise to an apparent higher and an apparent lower mass complex within an approximate mass of one TonB plus one FepA (∼ 116 kDa). [Although it has a calculated molecular mass of 26 kDa, TonB has an apparent mass of 36 kDa on SDS gels because 17% of its residues are prolines (41)]. Each of the two forms likely represented two different conformations made by the same complex. First, because only a single unoxidized Cys exists in FepA [because its two native Cys residues C487 and C494 are always oxidized (52)] and in the TonB Cys substitutions studied here, all of which carry the C18G substitution that removes the single native Cys. Second because the complexes are also detected with anti-FepA polyclonal antibodies, ruling out participation of a different protein (data not shown). And third, because similar doublets are also seen with TonB-BtuB disulfide-linked complexes *in vivo* (4). That disulfide-linked complexes can have different apparent masses depending on conformations of the participants is exemplified by the three different disulfide-linked TonB homodimers that result from a single TonB Cys substitution and demonstrably occur on non-reducing SDS polyacrylamide gels (17).

### Other FepA Cys substitutions assessed include the cork and periplasmic turns between β-strands of the barrel

Interaction of the TonB Cys substitutions with several additional FepA Cys substitutions other than the TonB box was assayed to identify potential additional sites of interaction by both the TonB amphipathic helix and Q160. In the mechanically weak domain of the FepA cork, L23, S29, T32, A33, D34, and E35 are periplasmically accessible in the crystal structure. Residues A42, S46, and G54 are buried; T51 is partially buried. L85 is in the mechanically recalcitrant domain and is partially buried [Fig. 9]. FepA residue T32, semi-conserved across TBDTs (53), and the less-well-conserved A33 were included because they bind to TonB at unknown sites and respond differentially to the presence and absence of ligand in FepA photocrosslinking studies *in vivo* (40). G54 exhibits a modest change in periplasmic exposure upon ligand binding (9).

**Figure 9:**
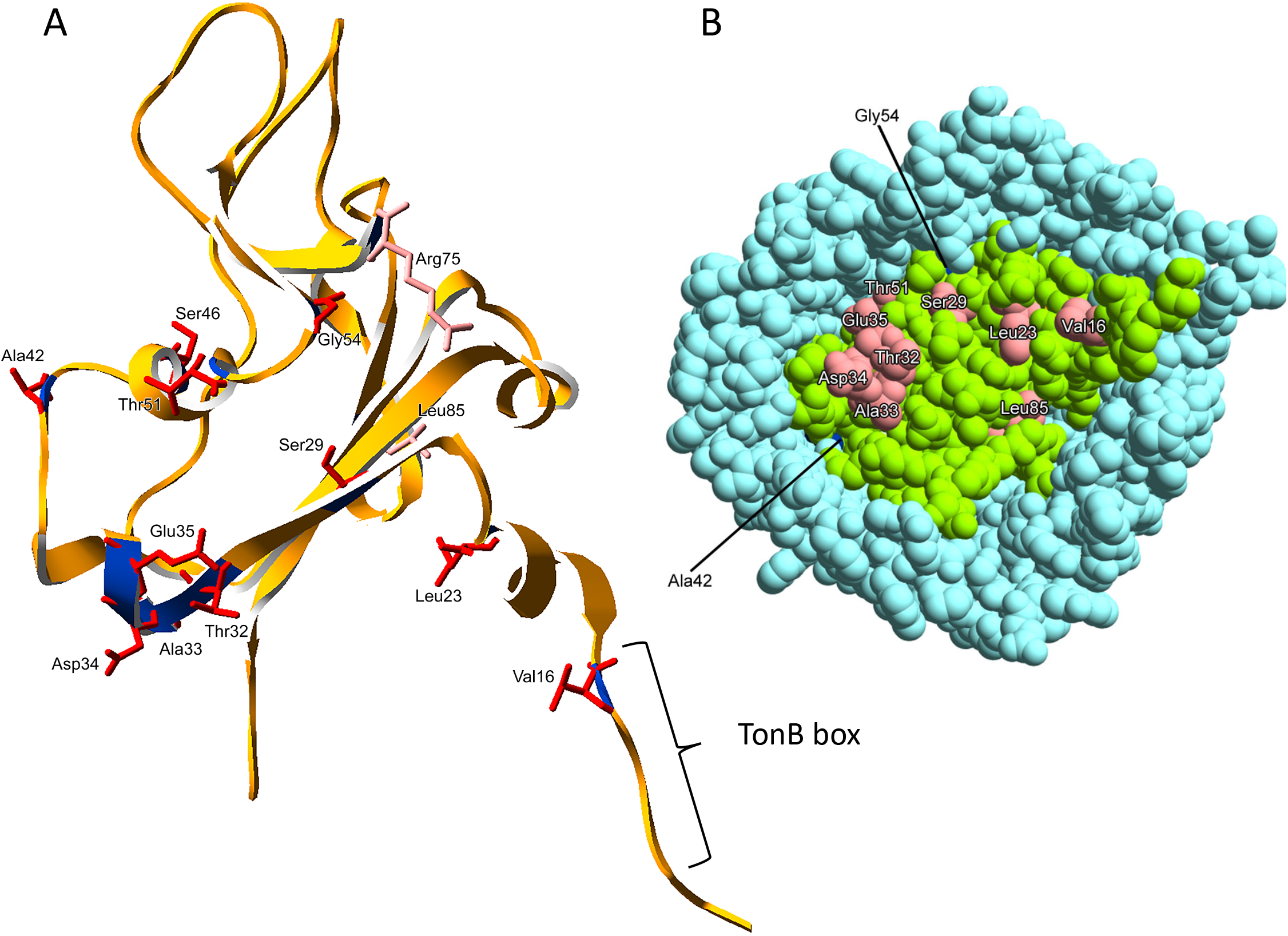
The FepA cork, residues 1-85. **(A)** Ribbon diagram of the FepA cork showing residues 1-85 in the mechanically weak segment of the FepA cork that were chosen for Cys substitution and *in vivo* disulfide crosslinking. The external surface of the cork is at the top, the periplasmic surface is at the bottom. The TonB box is indicated. Red residues formed crosslinks; pink residues did not. **(B)** Space filling model of the periplasmic surface of FepA, showing periplasmic accessibility of the residues labelled in (A). The FepA barrel is light blue; the FepA cork is light green; the periplasmically accessible residues tested are shown in pink. T51 is partially accessible; Gly54 and Ala42 (dark blue) are barely visible; Ser46 is completely buried. The crystal structure was solved by Buchanan et al. (60).

FepA residues in the mechanically recalcitrant segment of the FepA cork (V91, S92, S112, E120, V124, A131, V142, I145) were also evaluated to detect possible cork movements not observed previously [(12) Fig. 10]. Also evaluated were FepA R75, R126, E511, and E567, which form part of the “lock region”, with R75 and R126 in the recalcitrant domain of the FepA cork, and E511 and E567 positioned in the barrel (12, 14, 54). The lock region is proposed to be important for transport but not binding of ligand, with the positively and negatively charged residues forming a structure that keeps the cork bound to the barrel [(36, 55, 56).

**Figure 10.**
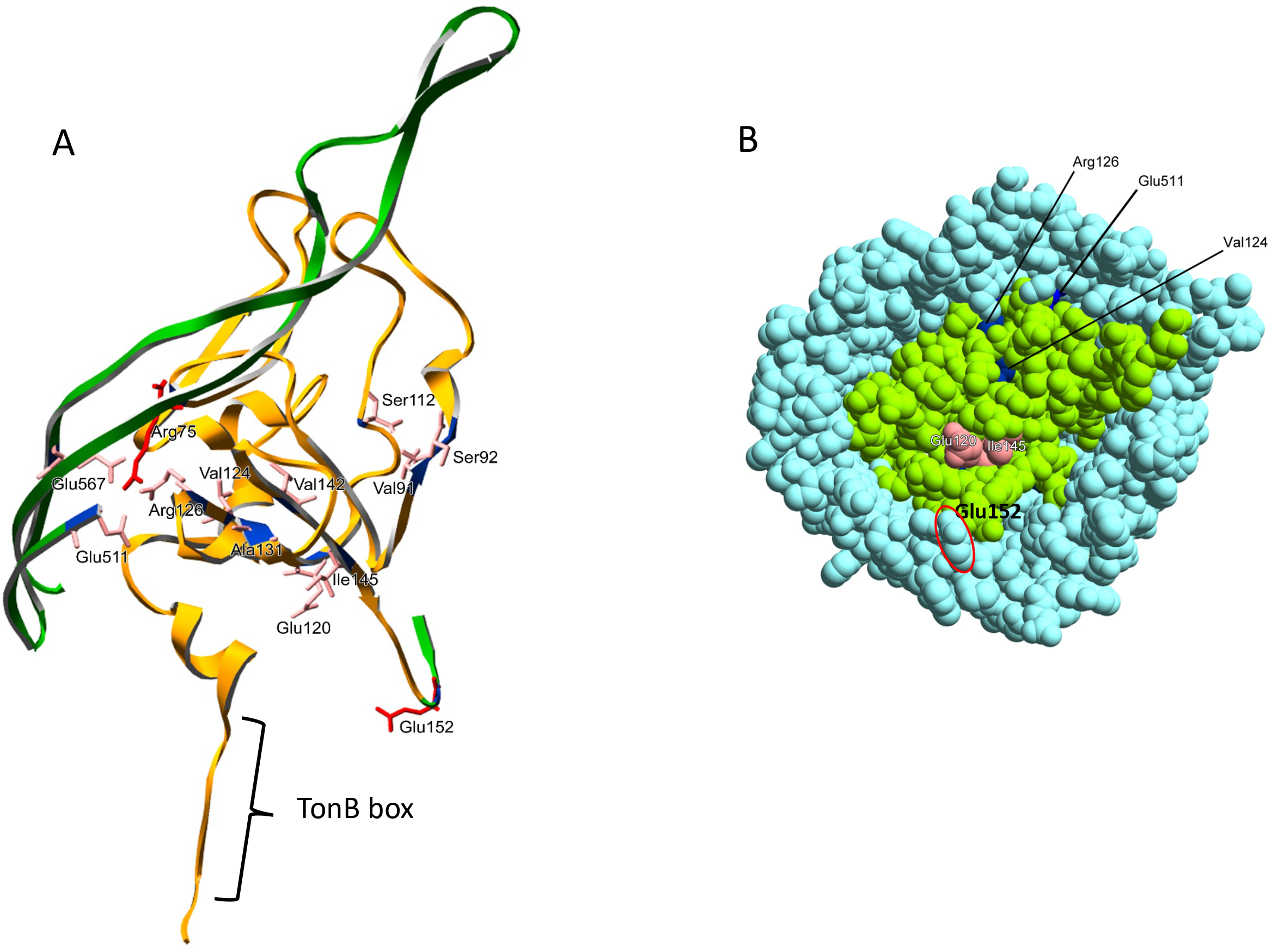
The FepA cork, residues 75-152. **(A)** Ribbon diagram of the FepA cork showing residues 75-152, a mechanically recalcitrant segment of the cork, that were chosen for Cys substitution and *in vivo* disulfide crosslinking. The external surface of the cork is at the top, the periplasmic surface is at the bottom. The TonB box is indicated. Red residues formed crosslinks; pink residues did not. Arg75 should be pink instead of red. Portions of the FepA barrel (dark green) show residues Glu511 and Glu567 which together with cork residues Arg75 and Arg126 form the lock region (54). **(B)** Space filling model of the periplasmic surface of FepA, showing periplasmic accessibility of the residues labelled in (A). The FepA barrel is blue; the FepA cork is light green; periplasmically accessible residues are shown in pink. Semi-accessible residues are dark blue. Glu 152 is circled in red. The crystal structure was solved by Buchanan et al. (60).

Contact with residues in periplasmic turns between β-strands of the FepA barrel, including the cork and barrel linker (D185, P243, D298, D356, D422, D455, D519, E576, D618, and D664; Fig. 11) were assessed, something that has not been investigated before for any TBDT. FepA residue E152, which marks the transition from cork to barrel and residue T722, the third residue from the carboxy terminus of FepA, were also included in the analysis.

**Figure 11.**
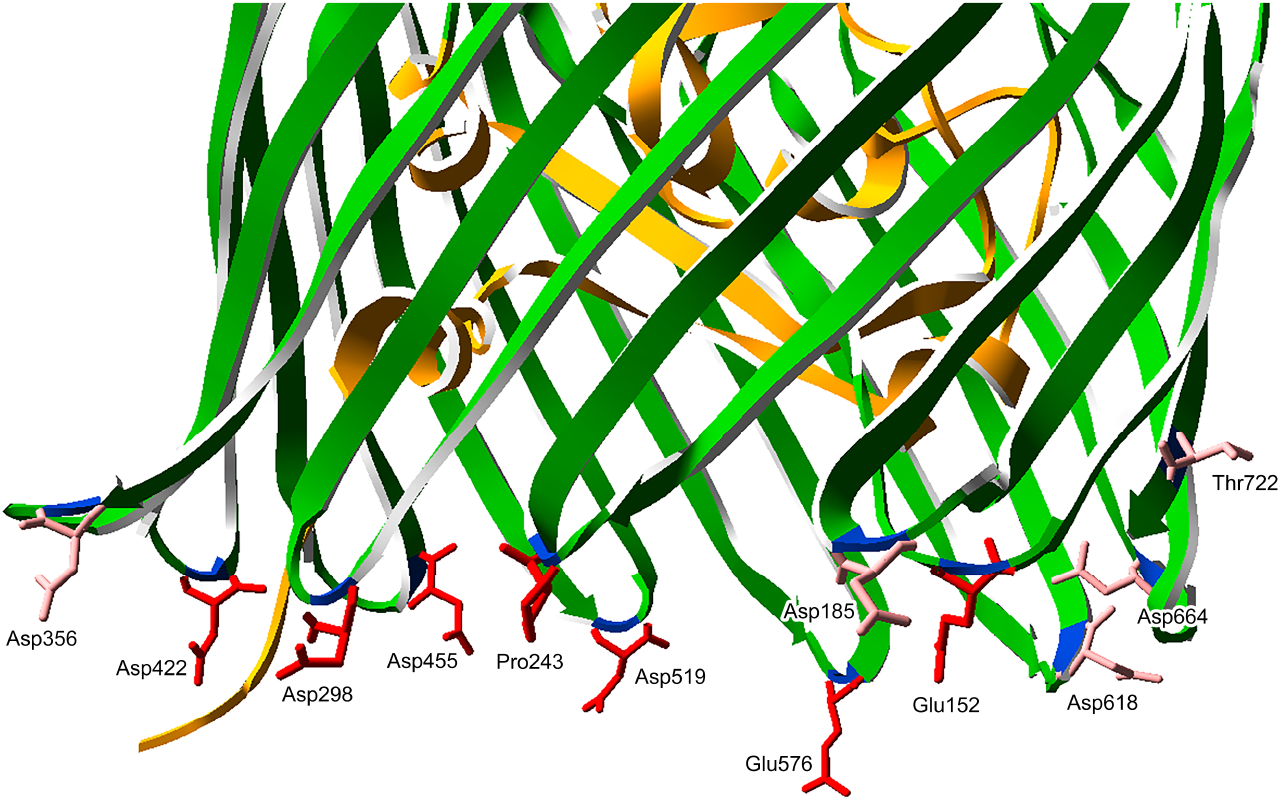
The FepA barrel β-strand turns. Ribbon diagram of the FepA β-strand turns chosen for Cys substitution and *in vivo* disulfide crosslinking. Beta strands of the barrel are colored dark green. Red residues formed crosslinks; pink residues did not. The crystal structure was solved by Buchanan et al. (60).

Overall, 35 additional Cys substitutions in FepA were tested for their ability to form disulfide crosslinks with TonB Cys substitutions.

### TonB amphipathic helix interactions extend into the mechanically weak region of the FepA cork

To define the boundaries of reactive residues in the TonB amphipathic helix, the set of core TonB amphipathic helix Cys substitutions was expanded to include M201C and R214C (Fig. 5). Each TonB Cys substitution was assayed pairwise at least twice in combination with the 35 additional FepA Cys substitutions outside the TonB box.

Together, these results indicate that only Cys residues in the mechanically weak region of the FepA cork interacted with TonB Cys substitutions, whereas mechanically recalcitrant cork region (substitutions R75C through I145C) and the two lock region residues in the barrel, E511C and E567C were essentially non-reactive (Fig. 12A, B). Several interactions also occurred with FepA periplasmic β-strand turns (Fig. 12C). All interactions involved the same reactive face of the amphipathic helix that interacted with the TonB box: R204C, N208C, and R212C. TonB M201C and R214C gave little to no interaction with any FepA Cys, confirming the boundaries of the core reactive residues. A key observation for a model to be described in the discussion was that none of the core amphipathic helix Cys residues (R204C, N208C, and R212C) interacted detectably with FepA Cys substitutions that were buried in the crystal structure (A42C, S46C, T51C, and G54C) (Figs. 9, 12A).

**Fig. 12.**
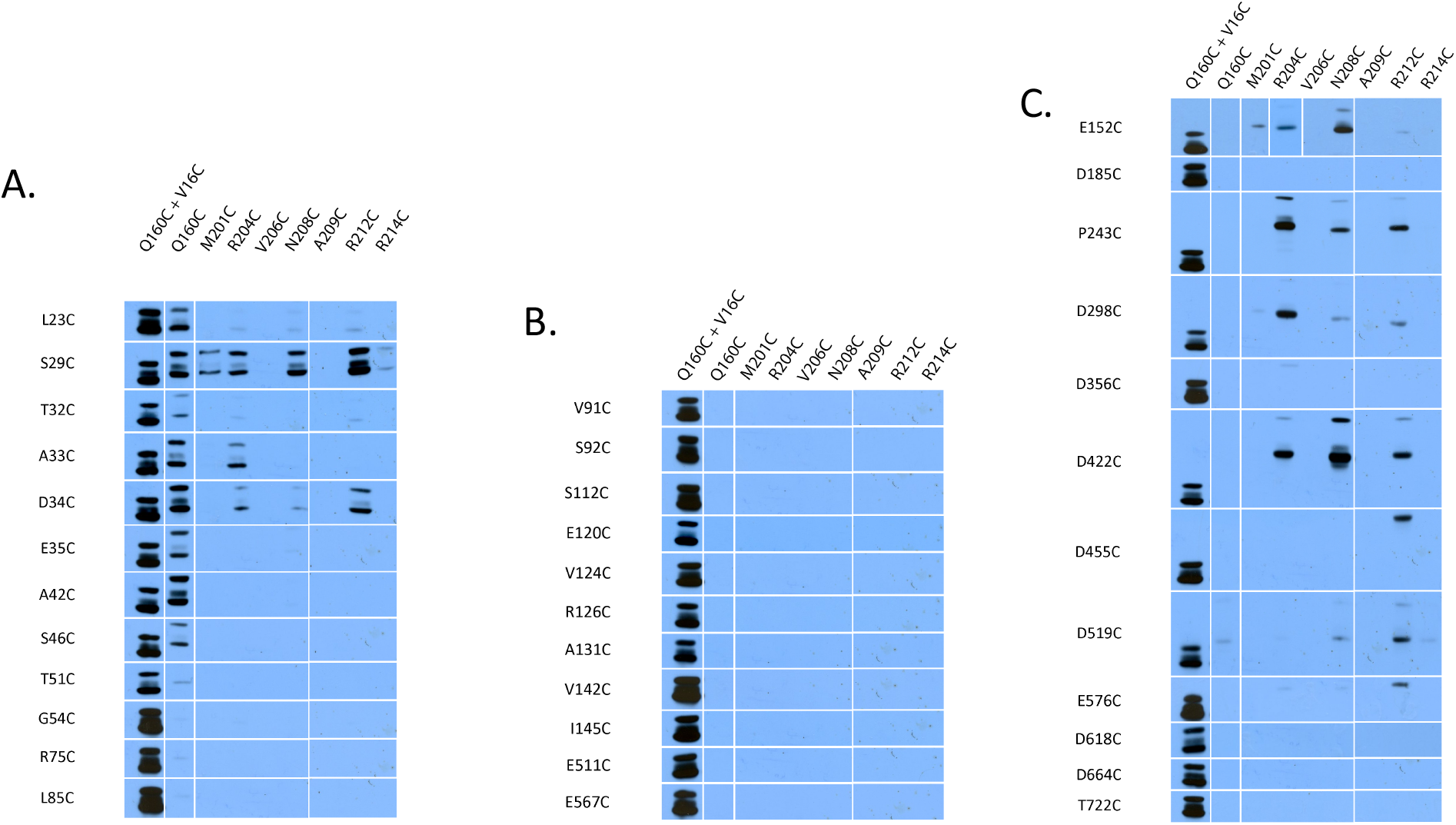
The composite comparison of *in vivo* TonB-FepA disulfide interactions identifies the interactive core of the TonB amphipathic helix as its hydrophilic face. **(A)** TonB Cys substitutions paired with FepA Cys substitutions from the mechanically weak region of the FepA cork (residues 1-70) as well as R75C and L85C from the transition between mechanically weak and mechanically recalcitrant domains. R75 is considered a part of the “lock region”. **(B)** TonB Cys substitutions paired with FepA Cys substitutions from the mechanically recalcitrant region of the FepA cork (residues 91-145) and the FepA “lock region” (residues R126, E511C and E567C). **(C)** TonB Cys substitutions paired with FepA Cys substitutions located in barrel β-strand turns as well as FepA E152C located at the transition between the FepA cork and barrel. TonB Cys substitutions are indicated across the top of the immunoblots. FepA Cys substitutions are indicated along the left side of each composite. TonB-FepA disulfide-crosslinked complexes were visualized in strain KP1491[W3110 Δ*fepA,* Δ(*tonB,P14*)*::kan*]. Composite immunoblots of non-reducing SDS polyacrylamide gels with anti-TonB monoclonal antibody are shown. Since not all experiments were performed on the same immunoblot, exposures for this composite summary were chosen based on matching the Q160C + V16C standards among immunoblots (left-most lanes in A, B, and C).

TonB R204C made several H20-dependent contacts with most of the periplasmically-accessible FepA cork Cys residues (L23C, S29C, T32C, A33C, and D34C, but not E35C), with S29C and A33C interactions being the most abundant (Fig. 13A).

**Figure 13:**
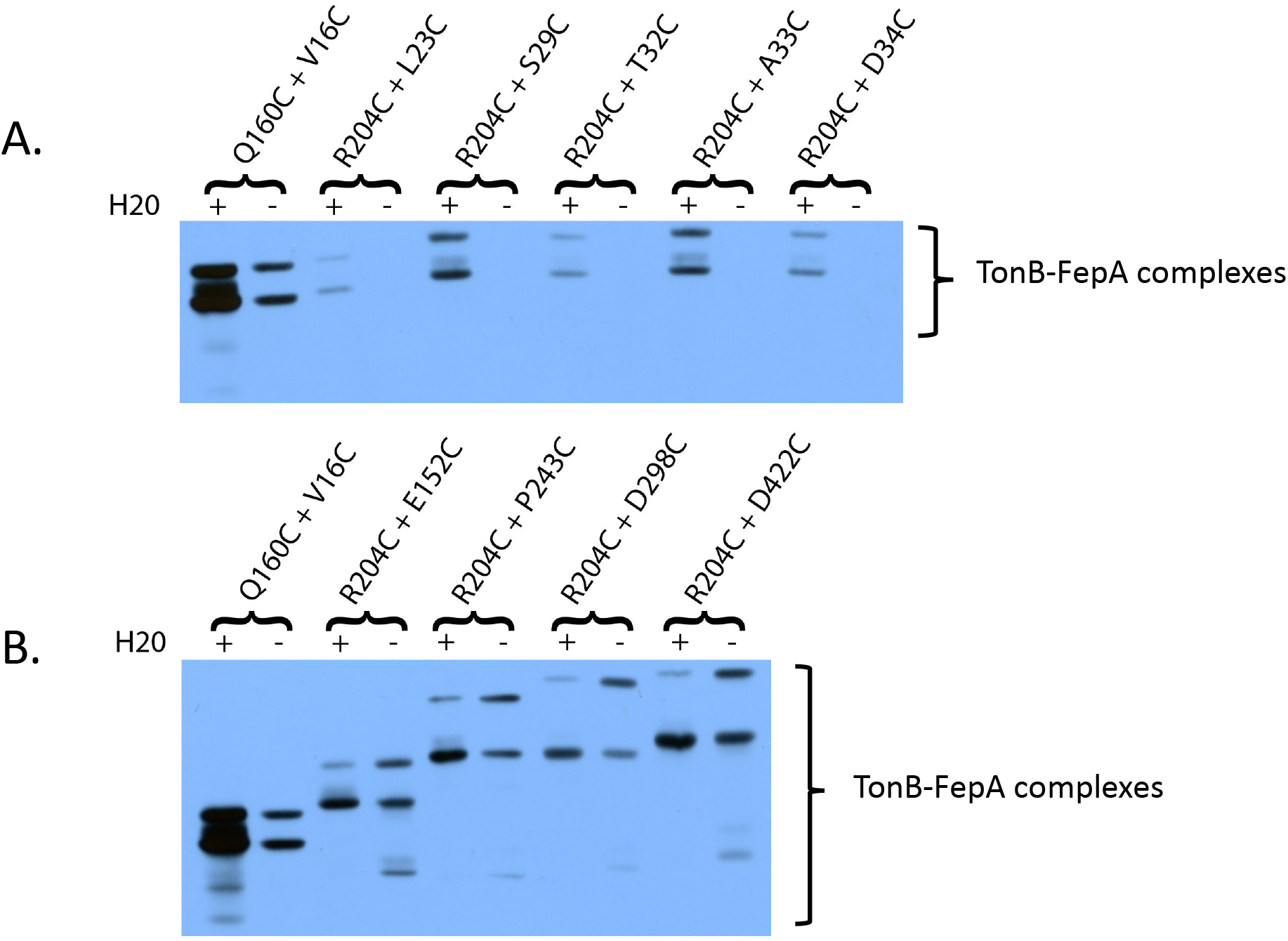
*In vivo*, TonB R204C makes both functionally important and functionally unimportant disulfide crosslinks with FepA Cys substitutions. TonB Cys substitution + FepA Cys substitution combinations are indicated at the top of each set of lanes. H20 (+) indicates the presence of the wild-type H20 allele in the TonB transmembrane domain. (-) indicates the presence of the inactivating H20A mutation. **(A)** The presence of the TonB H20A mutation significantly reduced disulfide crosslinking by FepA early cork Cys substitutions (depicted in Fig. 9). **B).** The H20A mutation has little effect on the overall abundance of R204C-mediated disulfide crosslinks at the cork-barrel interface, specifically FepA E152C (depicted in Fig. 10) or the barrel turns, where they occurred (depicted in Fig. 11). The TonB Q160C-FepA V16C pair was used as a standard for comparison of relative levels (far left lanes in A. and B.). TonB-FepA disulfide-crosslinked complexes were visualized in strain KP1491[W3110 Δ*fepA,* Δ(*tonB,P14*)*::kan*]. Immunoblots of non-reducing SDS polyacrylamide gels with anti-TonB monoclonal antibody are shown. Monomer TonB levels for all samples in these immunoblots were at or near chromosomal levels (data not shown).

The only residue in the cork region significantly contacted by TonB N208C was FepA S29C and it was an H20-dependent interaction (Fig. 14A).

**Figure 14:**
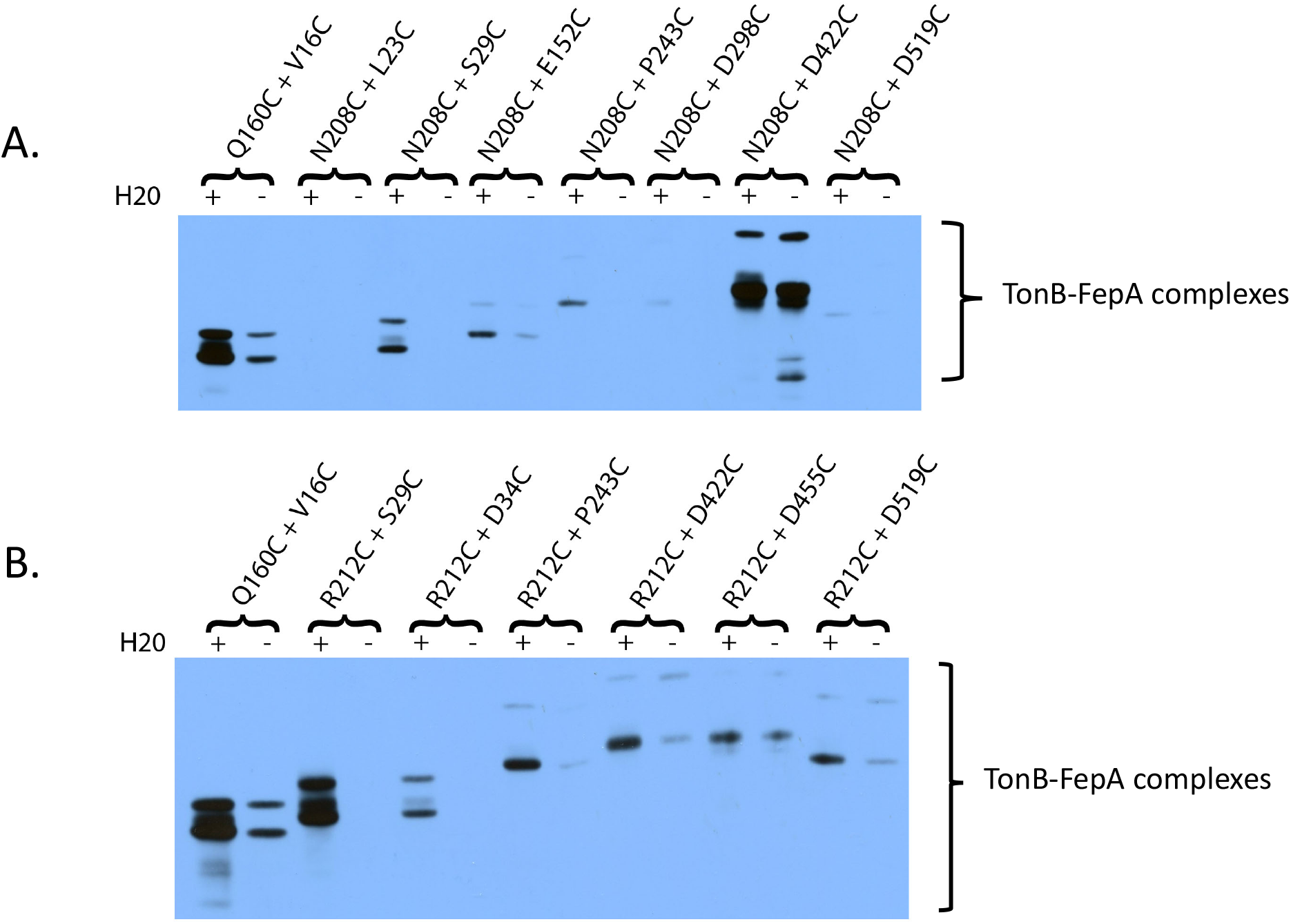
*In vivo*, TonB N208C and TonB R212C make both functionally important and functionally unimportant disulfide crosslinks with FepA Cys substitutions. TonB Cys substitution + FepA Cys substitution combinations are indicated at the top of each set of lanes. H20 (+) indicates the presence of the wild-type H20 allele in the TonB transmembrane domain. (-) indicates the presence of the inactivating H20A mutation. **(A)** The presence of the TonB H20A mutation generally reduced disulfide crosslinking by TonB N208C. The notable exception was the very abundant crosslink with D422C, located in a FepA barrel turn (depicted in Fig. 11). This interaction was impervious to the presence of the H20A mutation. **(B)** Most notably, TonB R212C makes a very abundant, H20-specific, crosslink with FepA S29C (depicted in Fig. 9). It also makes H20-specific complexes with some of the FepA Cys substitutions in the barrel turns (depicted in Fig. 11). The TonB Q160C-FepA V16C pair was used as a standard (far left lanes in A. and B.). TonB-FepA disulfide-crosslinked complexes were visualized in strain KP1491[W3110 Δ*fepA,* Δ(*tonB,P14*)*::kan*]. Immunoblots of non-reducing SDS polyacrylamide gels with anti-TonB monoclonal antibody are shown. Monomer TonB levels for all samples in these immunoblots were at or near chromosomal levels (data not shown).

TonB R212C made the highest degree of H20-dependent contacts with FepA S29C—as high as the contacts between TonB Q160C and the FepA TonB box residue V16C, which were the highest we observed (Figs. 7, 14B).

FepA S29C appeared to be a hot spot since R204C, N208C, and R212C as well as the boundary-defining TonB M201C and R214C made complexes with it. FepA S29 is located close to the center of the first β-strand of the cork. It is within the region through which TonB could mechanically pull to unravel the cork as suggested before for FhuA (20), possibly as the site of *in vivo* TonB interaction at the as-yet-to-be-identified non-TonB box site prior to TonB box exposure (12).

### The TonB amphipathic helix interacts with periplasmic FepA β-strand barrel turns

*In vivo*, TonB interacts with FepA at one or more sites before it interacts with the FepA TonB box (12). Instead of FepA S29, the periplasmic β-strand turns of TBDTs could constitute such sites.

The last 150 residues of the TonB periplasmic domain, within which the amphipathic helix resides, have a calculated pI of 10.4 (41). It seemed logical that this TonB domain would be most likely to interact with the negatively charged residues of FepA barrel turns, where there appears to be one in nearly every turn (except Pro243 in turn 2), including Glu152 in the linker region between cork and β-strand 1. Such interactions would likely be mediated through additional residues of TonB and FepA that would bring the now Cys-substituted residues into proximity (Fig. 11).

All of the FepA Cys substitutions in β-strand turns exhibited the ability to support fully wild-type levels of ^55^Fe-enterochelin transport except FepA E152C and T722C, where the levels dropped to ∼ 60% (Table 1). Two Cys substitutions in the lock region of the barrel, E511C and E567C, were also assayed. FepA E511C had little effect on FepA activity, and E567C reduced activity to 65% of wild-type. Cys substitutions in the cork components of the lock region, R75C and R126C, reduced transport to ∼40% of wild-type levels but did not eliminate it. Thus, neither the charged residues in the periplasmic β-strand turns nor the lock region residues were individually essential for FepA function.

Several sites of interaction between the TonB amphipathic helix residues R204C, N208C, and R212C and the periplasmic FepA β-strand barrel turns are summarized in Fig. 12C.

For TonB R204C, two different forms of the complex were observed, as was also seen with the FepA cork Cys substitutions (Fig. 13B). In contrast to interactions with the FepA cork, the apparent masses of the complexes with E152C, P243C, D298C, and D422C shifted to significantly higher values, highest in the case of the latter three for unknown reasons. In contrast to interactions with the cork, the β-strand barrel complexes were largely insensitive to the presence of TonB H20A. Overall intensities of the complexes remained similar with both TonB H20 and TonB H20A versions of R204C, but in the case of TonB H20A, the complexes appeared to slightly shift their abundance to the higher mass complex— again for unknown reasons.

For TonB N208C, weak interactions were detected with FepA E152C, P243C, D298C, and D519C, each of which was H20-dependent (Fig. 14A). In addition, TonB N208C made a very high level of contacts with D422C whether TonB H20 or H20A was present. TonB R212C made a moderately high level of H20-sensitive contacts with P243C, D422C, and D519C, whereas the contact with D455C was largely H20-insensitive (Fig. 14B). Thus, both functional and non-functional TonB amphipathic helix contacts occurred with the FepA β-strand turns.

### TonB Q160 interactions include buried cork residues

Previous studies of TonB Q160C with TBDTs have been confined to the TonB box. Since the TonB amphipathic helix made FepA contacts outside the TonB box, we wanted to explore the possibility that Q160 did so as well.

Surprisingly, TonB Q160C interacted more widely than the amphipathic helix did with Cys substitutions throughout the mechanistically weak part of the FepA cork (Figs. 15A, B). Disulfide-linked heterodimers were observed between TonB Q160C and FepA V16C, L23C, S29C, T32, A33C, D34C, E35C, A42C, S46C and T51C, with the highest degree of interactions occurring with D34C and S42C. The presence of the H20A mutation greatly diminished, and in most cases eliminated, the crosslinking detected, suggesting that when TonB was inactive, the ability of Q160C to make contacts within the mechanically weak domain of the FepA cork was entirely prevented, unlike the interaction with the FepA TonB box (Fig. 7A; 15C)

**Figure 15:**
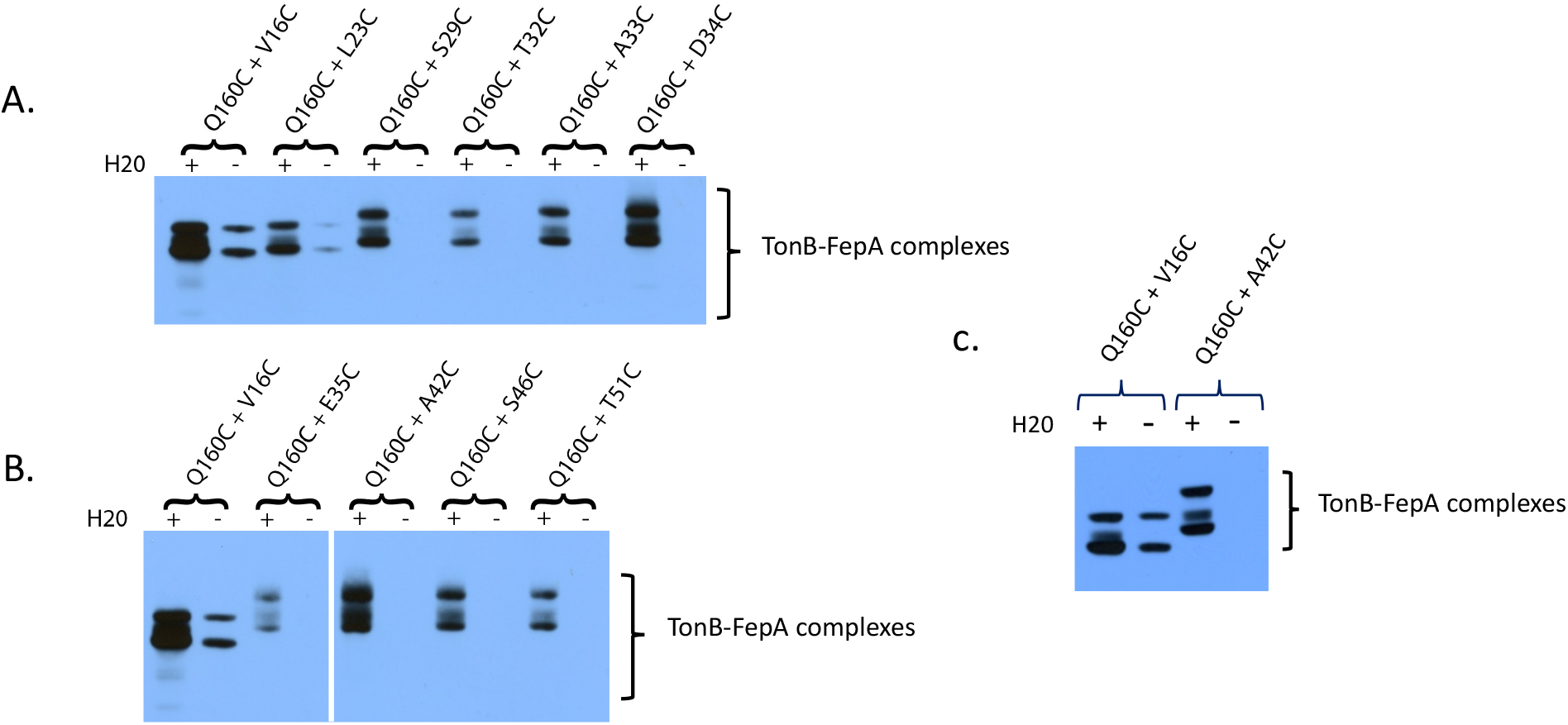
*In vivo*, TonB Q160C makes several functionally important disulfide crosslinks within the mechanically weak segment of the FepA cork, including with buried residues (depicted in Fig. 9). TonB Cys substitution + FepA Cys substitution combinations are indicated at the top of each lane. H20 (+) indicates the presence of the wild-type H20 allele in the TonB transmembrane domain. (-) indicates the presence of the inactivating H20A mutation. **(A)** TonB Q160C made complexes with several FepA substitutions, all of which were prevented by the TonB H20A mutation. Notably the complex with FepA D34 was of nearly equal abundance to the standard TonB Q160C-FepA V16C pair; unlike that standard, it was entirely prevented by the presence of the TonB H20A mutation. **(B)** TonB Q160C makes a complex with buried residue FepA A42C of nearly equal abundance to the standard TonB Q160C-FepA V16C pair; it was entirely prevented by the presence of the TonB H20A mutation. For this composite immunoblot, the exposure on the right was chosen based on matching it to the same intensity as the Q160 + V16C standard shown in the left panel. **(C)** Direct comparison of Q160C complexes with FepA V16C and FepA A42C on the same immunoblot and with a shorter exposure. TonB-FepA disulfide-crosslinked complexes were visualized in strain KP1491[W3110 Δ*fepA,* Δ(*tonB,P14*)*::kan*]. Immunoblots of non-reducing SDS polyacrylamide gels with anti-TonB monoclonal antibody are shown. Monomer TonB levels for all samples in these immunoblots were at or near chromosomal levels (data not shown).

The disulfide crosslinking between TonB Q160C and FepA A42C or S46C was notable because A42 and S46 are buried in the FepA crystal structure. TonB Q160C crosslinks with FepA A42C were as abundant as those between the standard TonB Q160C-FepAV16C pair (Figs. 9 and 15C). FepA A42 and S46 are positioned in a plane approximately mid-way between top and bottom of the cork. FepA A42 is on the interface with the FepA barrel. FepA S46 is entirely buried within the cork (Fig. 9). These key observations are incorporated into a model in the discussion.

TonB Q160C did not interact abundantly with any Cys substitutions more carboxy-terminal than FepA T51C (Fig. 12). Consistent with cork movement, weak H20-specific interactions with FepA G54C were also observed on long exposures (Fig. 16, lane 5). Ma et al. previously observed modest periplasmic exposure of FepA G54C in the presence of enterochelin (9). With the exception of scarcely detectable interactions with D519C in a barrel turn, no interactions of TonB Q160C with any of the remaining FepA Cys substitutions from V91C through I145C (the mechanically recalcitrant domain), the lock region or the barrel turns were detected no matter how long the exposures were (Figs. 12B and 12C; data not shown). These observations form an important part of the model presented in the discussion.

**Figure 16.**
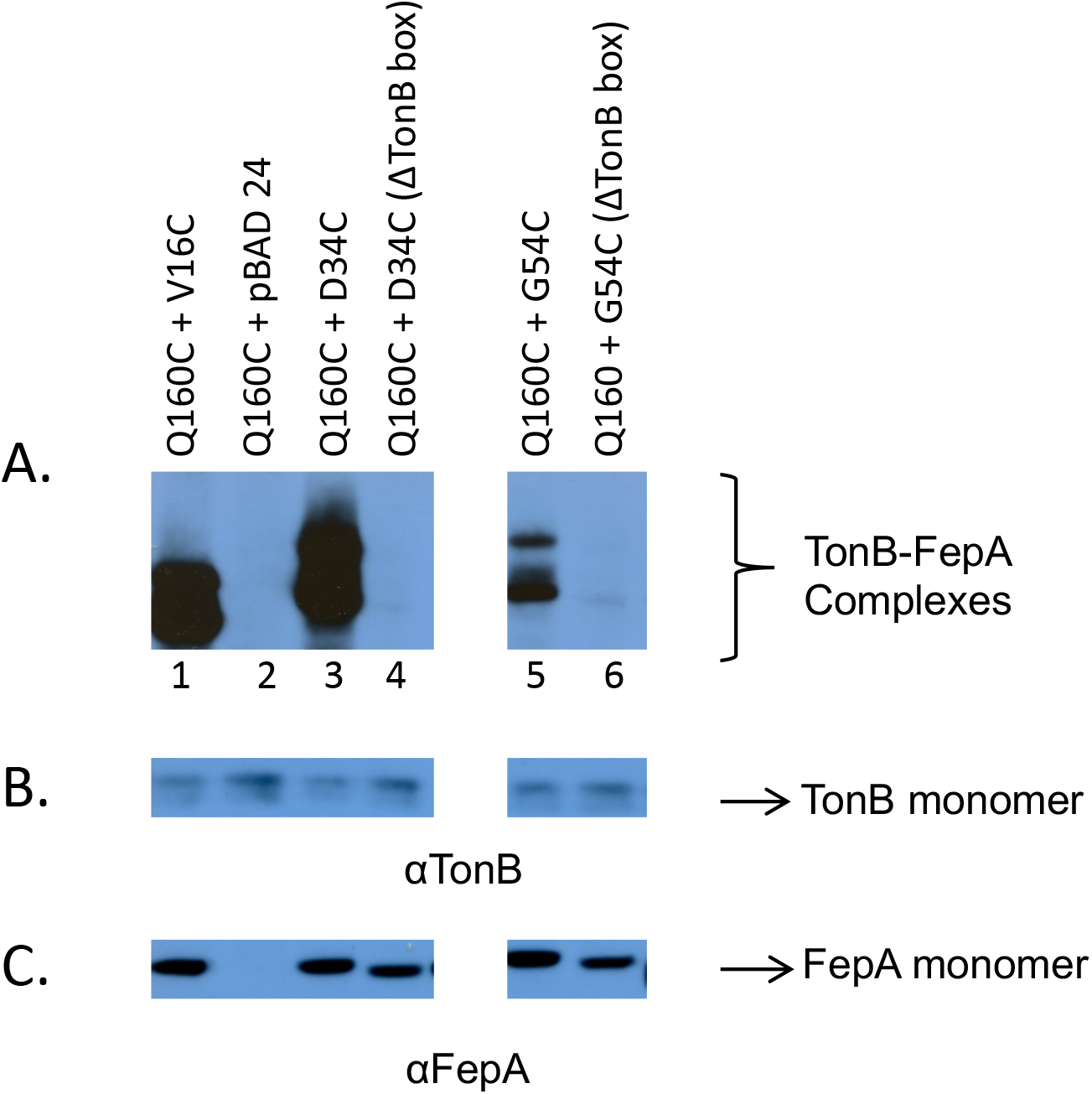
Deletion of the FepA TonB box prevents *in vivo* disulfide complex formation by TonB Q160C. **(A)** TonB-FepA disulfide-crosslinked complexes visualized in strain KP1491[W3110 Δ*fepA,* Δ(*tonB,P14*)*::kan*] by immunoblots of non-reducing SDS polyacrylamide gels with anti-TonB monoclonal antibody. pBAD24 is the vector into which *fepA* variants were cloned. **(B)** Monomer levels of TonB from the same samples as A), visualized by anti-TonB monoclonal antibody. **(C)** Monomer levels of FepA from same samples as A.), visualized by anti-FepA polyclonal antibody. All lanes are from the same immunoblot with a center lane masked.

### Deletion of the FepA TonB box prevents Q160C FepA cork interactions

Prior to the experiments above, the interaction of TonB Q160 with TBDT sites other than their TonB boxes had not been tested. Since several additional interactions were identified, we attempted to determine an order of events by analyzing the effect of a FepA TonB box deletion on complex formation with the FepA Cys substitution D34C with which TonB Q160C interacts with as abundantly as it does the FepA TonB box, and G54C, which is buried and where the interaction is weaker. In both cases, deletion of the FepA TonB box (residues 12-17) essentially prevented the interaction (Fig. 16A, lanes 4 and 6). This finding suggested that prior contact by unspecified TonB residues with the FepA TonB box was required for TonB Q160C to interact with FepA cork residues beyond the TonB box.

### TonB F202A, W213A lacks ExbD contact and inhibits the interaction of TonB Q160C with the FepA TonB box

In TonB, the F202A and W213A mutations boundary the core of the amphipathic helix and, in combination, completely inactivate it (29). TonB F202A, W213A was used as a tool to better understand parameters of the interaction between TonB Q160C and FepA V16C in the TonB box.

At the time when we discovered the inactivity of TonB F202A, W213A, the TonB-ExbD interaction captured by *in vivo* formaldehyde crosslinking of TonB had yet to be identified, however we knew that such double Ala mutations in the carboxy terminus did not prevent formation of the disulfide-linked TonB triplet homodimers (17, 29). In this study, the effect of the F202A, W213A mutation as well as another double Ala mutation--F202A, Y215A--was to prevent formation of the Stage III TonB-ExbD formaldehyde crosslinked heterodimer (18, 31). It was particularly telling that even when the TonB double Ala mutants were overexpressed, there was still no formation of the TonB-ExbD heterodimer, which appears to play a key role in configuring TonB for successful energy transduction to FepA (Fig. 17; Fig. 2, Stage III TonB-ExbD heterodimers).

**Figure 17.**
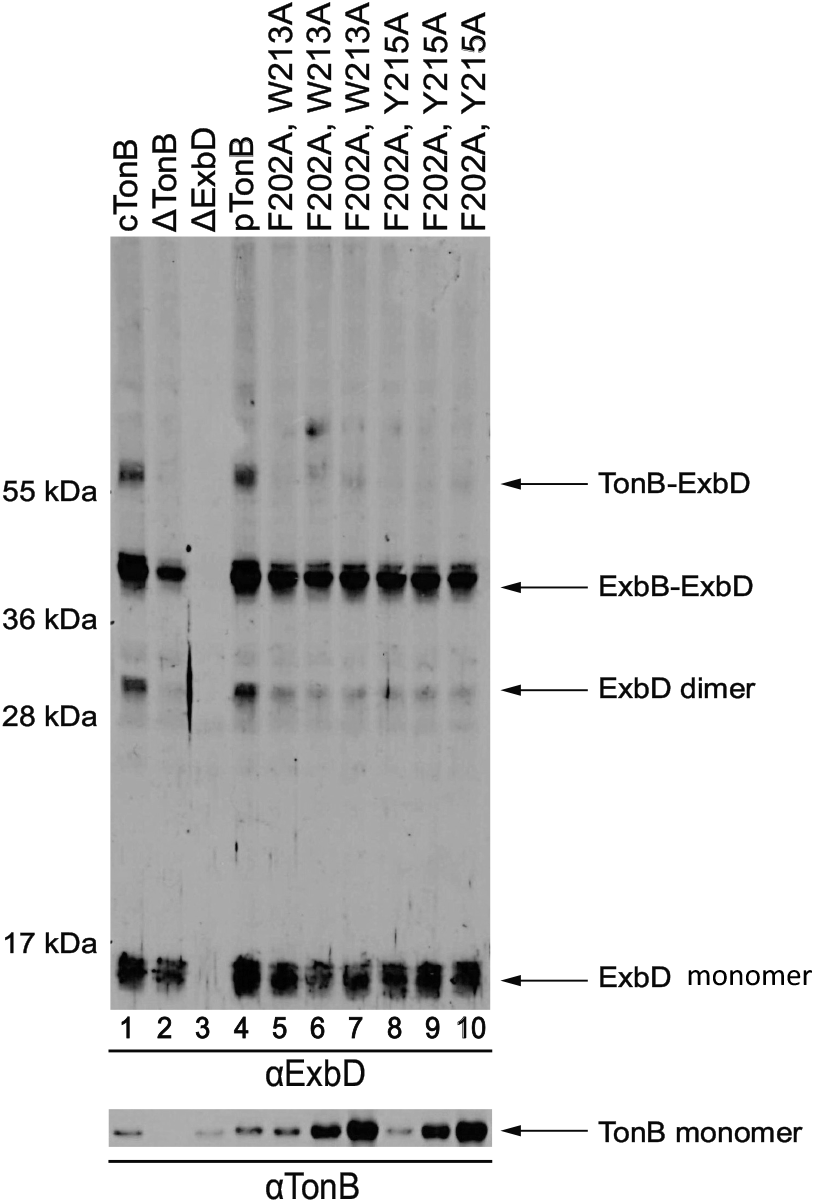
TonB F202A, W213A does not form the TonB-ExbD heterodimer. **Upper:** Immunoblot with anti-ExbD antibodies of formaldehyde crosslinked samples. Lane 1, W3110 expressing chromosomally encoded TonB (cTonB); Lane 2, KP1344 [W3110 Δ(*tonB, P14)::blaM*]; Lane 3 RA1021 (W3110 Δ*exbD*); Lane 4, plasmid-encoded TonB (pKP442) with 0.001% arabinose; Lane 5, pKP531 (pKP442 TonB with F202A, W213A double mutations) with 0.002% arabinose; Lane 6, pKP531 with 0.005% arabinose; Lane 7, pKP531 with 0.01% arabinose; Lane 8, pKP532 (pKP442 TonB with F202A, W215A double mutations) with 0.002% arabinose; Lane 9, pKP532 with 0.005% arabinose; Lane 10, pKP532 with 0.01% arabinose. **Lower:** Corresponding steady state levels of TonB from the samples above are shown. For pKP531 and pKP532, note the increase in TonB expression with increasing addition of the inducer, arabinose.

When F202 and W213 are mutated individually, they support intermediate and assay-specific levels of TonB activity (29). Consistent with that, the TonB Q160C, F202A substitution was still able to form the Q160C-V16C complex (Fig. lane 4). Although the presence of the F202A, W213A double mutations rendered TonB proteolytically unstable (Fig. 18, lane 5), as seen previously (29), it was possible to increase the exposure time of the immunoblot to the point where the level of monomer (Fig. 18, lane 6) was slightly greater than the level seen for the TonB Q160C and its F202A derivative (Fig. 18, lanes 3 and 4). In the longer exposure, it was clear that the ability of the TonB Q160C, F202A, W213A to form complexes with FepA V16C was significantly diverted away from the FepA TonB. Instead, the absence of a functional TonB carboxy terminus led Q160C to form three complexes, too small to be complexes with FepA. The location and spacing of the complexes were reminiscent of disulfide-linked TonB triplet homodimers that represent three different conformations of the TonB carboxy terminus *in vivo* (17).

**Figure 18.**
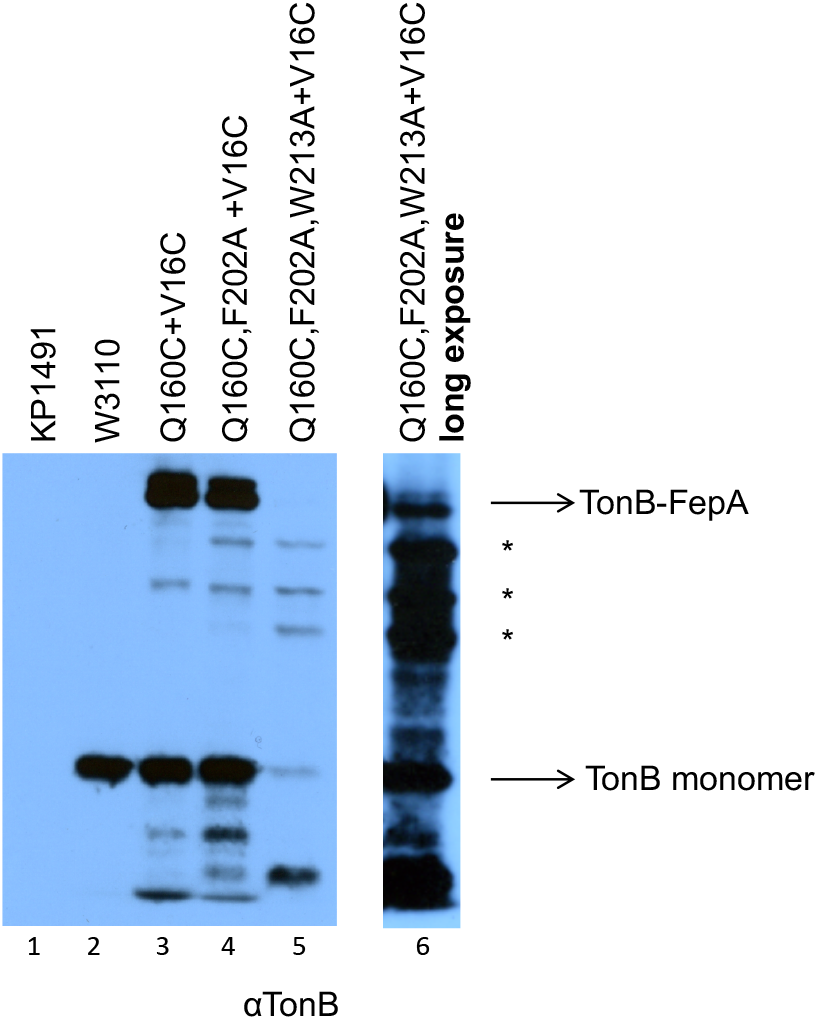
Inactivation of the TonB carboxy terminus appears to divert TonB Q160C down a path to triplet homodimer formation. Lane 1, KP1491 [W3110 Δ*fepA,* Δ(*tonB,P14*)*::kan*]: the parent strain used in all these studies. Lane 2, W3110: the wild-type strain showing the steady state level of chromosomally encoded TonB. Lane 3, TonB Q160C in combination with FepA V16C. Lane 4, TonB Q160C with the additional F202A substitution in combination with FepA V16C. Lane 5, TonB Q160C in combination with additional F202A and W213A substitutions in combination with FepA V16C. Lane 6, long exposure of lane 5 to reveal relative levels of TonB-FepA and triplet homodimer complexes. Positions of TonB monomer and TonB-FepA complexes are indicated on the right. Potential triplet homodimers are indicated by asterisks (*). Immunoblots of non-reducing SDS polyacrylamide gels developed with monoclonal anti-TonB antibody are shown.

## DISCUSSION

While TonB remains anchored in the cytoplasmic membrane by its amino terminal transmembrane domain, the periplasmically localized TonB carboxy terminus binds transiently and cyclically to TonB-dependent transporters (TBDTs) in the outer membranes of Gram-negative bacteria to transduce cytoplasmic membrane protonmotive force energy required for the active transport of ligands (8). The *in vivo* molecular mechanism is not well understood. Previously, the only known *in vivo* interaction between TonB and TBDTs involved the region of TonB Q160 and the BtuB and FecA TonB boxes (4, 16, 50). Because our earlier studies had suggested additional but unknown sites are involved *in vivo*, we searched for site-specific interactions between TonB and FepA, the TBDT that transports the sole siderophore synthesized and secreted by *E. coli* (12). TonB sites focused on TonB Q160 and the TonB carboxy terminal amphipathic helix and their potential interactions with sites in the FepA TonB box, in both mechanically weak and mechanically recalcitrant domains of the FepA cork, and in FepA β-strand barrel turns. It is important to study TonB interactions with TBDTs in their native environment, where ExbB, ExbD and the protonmotive force of the cytoplasmic membrane are all present (57).

### The TonB carboxy terminal amphipathic helix is essential for the energy transduction cycle

TonB encoded from an amber mutation at codon 175 is inactive, demonstrating that, although TonB Q160 has been the only site established to interact with TBDTs, it is not sufficient for activity, and indicating that some aspect of the last 65 TonB residues is essential (37). Within those last 65 residues, we focused on the role of a sole amphipathic helix in the TonB carboxy terminus (residues 199-216).

We found that the TonB amphipathic helix was essential either by deleting it or by frameshifting it to maintain overall residue continuity. While this confirmed the importance of the region, those results could also reflect an inability to form TonB-TonB homodimers or TonB-ExbD heterodimers, both of which are important for the energy transduction cycle (27, 31, 34, 35). We therefore asked if the TonB amphipathic helix interacted directly with the only known essential region of the TBDT, FepA, its TonB box.

Based on the *in vivo* disulfide crosslinking experiments, amphipathic helix residues R204C, N208C and R212C defined a hydrophilic face that interacted with Cys substitutions throughout the FepA TonB box, whereas the two residues on the hydrophobic face, V206C and A209C did not interact with FepA. TonB R204C was previously shown to be solvent exposed at some point in the energy transduction cycle, consistent with localization on the hydrophilic face (58). Because these *in vivo* interactions were prevented by the presence of the inactivating TonB H20A mutation, they comprised a set of novel, functional, and specific interactions that have been identified between TonB and a TBDT for the first time.

The amphipathic helix interaction with the TonB box is absent from solved co-crystal structures of the TonB carboxy terminus (∼ residues 152-235) with TBDTs BtuB and FhuA. The sidechains of the hydrophilic face residues R204C, N208C, and R212C are oriented towards the TBDTs and distant from the TonB boxes in both structures (20, 45), demonstrating either the difference between *in vivo* “energized” and inactive TonB, or differences between TBDTs.

Now that an essential TonB component has been identified that interacts with an essential FepA component (TonB box), it is tempting to speculate that the TonB amphipathic helix holds the entire key to the TonB energy transduction cycle for *E. coli.* As a result of this study and previous work, residues within the TonB amphipathic helix have now been recognized to participate in sequential interactions with three different proteins—TonB with itself, with ExbD and, here, with FepA [(24, 59); Fig.2]. The amphipathic helix sequences are 55% conserved (72% if E/D, R/K, and W/F substitutions are considered equivalent) amongst enteric bacteria, but barely conserved with *Pseudomonas putida tonB*, which does not complement an *E. coli tonB* mutation (24, 59).

Like wild-type TonB, TonB H20A formaldehyde-crosslinks with FepA *in vivo*, but does not transmit energy to it (18, 28, 44). Formation of disulfide crosslinks and their elimination by the inclusion of the TonB H20A mutation was a clear indication that they represented functional interactions. While the lack of disulfide-linked complexes for some pairs likely represented lack of interactions, it could also be that the interactions could not be trapped due to due to misorientation of the thiol side chains or because the interactions were too transient. It is also likely that other important *in vivo* regions of interactions remain to be discovered.

### A model: does the FepA cork “fish” for the TonB Q160 region?

On the face of it, the amphipathic helix region (residues 199-216) of TonB is more logical than TonB Q160 as a site of initial contact with a TBDT TonB box because it is theoretically somewhat closer to the outer membrane. This is especially so since ∼ 100 Å of TonB’s reach across the periplasmic space is due to the proline-rich domain (residues 70-102), which can be deleted without eliminating TonB activity [Fig. 4; (24, 39)]. In addition, the amphipathic helix is essential, whereas TonB Q160 and the region encompassing it (residues R158-Q162) can be deleted without inactivating TonB (26). Why this region has been a source of second site suppressors for inactive TBDT TonB box mutations remains a mystery (48, 49).

Because it is not essential, the wide range of *in vivo* contacts made by TonB Q160C was surprising, encompassing not only the FepA TonB box seen previously for other *E. coli* TBDTs but also, for the first time, residues throughout the region of the mechanically weak domain of the FepA cork. Most importantly, two of the interacting residues (A42, and S46) are buried within in the FepA cork (60). In contrast to the partially H20-senstive Q160C interactions with the FepA TonB box, both of these contacts were completely prevented if the TonB Q160C also carried the inactivating H20A mutation, indicating that they fully represented the action of functional TonB. The FepA region of TonB Q160C interaction ended just prior to the beginning of the mechanically recalcitrant domain of the FepA cork.

TonB Q160C interacted as abundantly with FepA A42C as it did with the FepA V16C standard within the TonB box, which was the most abundant interaction seen in this entire study. Because FepA A42C is buried, this strongly suggested that enough of the cork domain entered the periplasm to expose A42C, along with S46C, to interaction with TonB Q160C. Our previous discovery of 20-to-25-fold increases in periplasmic accessibility of the buried cork residues A42C, S46C (and the partially buried T51C) in response to the addition of colicin B ligand *in vivo* validates the finding here that at some point in the energy transduction cycle, these residues become available for interaction with TonB Q160C and likely neighboring residues (4, 12). In striking contrast, none of the amphipathic helix Cys substitutions interacted with FepA Cys substitutions that were buried within the cork, with targets limited to periplasmically accessible Cys residues.

The overall non-reactivity of the mechanically recalcitrant domain of FepA in this *in vivo* study also validated our earlier finding of its resistance to being periplasmically labeled in the presence of a large (∼55 kDa) ligand, colicin B. In that study, two distinct possibilities were proposed for the mechanically recalcitrant domain: first that it did not move and second that it moved but was blocked from being labeled by the presence of another protein (12). Our studies here did not exclude either possibility. For the first possibility to be true, colicin B would need to denature on its way through a small opening, which, given its size and structure as a dumbbell that fills the FepA lumen, seems unlikely (61). Nonetheless, such denaturation has been observed for the amino-terminal domain of colicin pyoS2 of *P. aeruginosa* through its TBDT FpvA *in vivo* (62). Furthermore, the significantly smaller (∼29 kDa) colicin M, which parasitizes the TBDT FhuA, uses a chaperone to fold it in the periplasm where it inhibits peptidoglycan formation (63). A mechanically recalcitrant domain for the TBDT BtuB has been characterized *in vitro* (14).

While it is not possible to definitively turn static data into a dynamic model, these results suggested a possible sequence of events where the TonB amphipathic helix binds to the FepA TonB box, which moves the FepA cork sufficiently that its mechanically weak domain extends into the periplasm. Previously buried cork residues are thus able to “fish” for interactions with various sites on TonB, among which we captured the TonBQ160C interaction. It is notable that the hydrophilic face of the amphipathic helix contacted multiple residues in the β-strand turns of the FepA barrel, whereas TonBQ160C could contact none of them, consistent with the idea that it does not reach that far across the periplasmic space. These results may also explain why the Q160 region of TonB is not essential—it is secondary and incidental to the action of the TonB amphipathic helix at the FepA TonB box.

Consistent with this model, without the FepA TonB box present for the amphipathic helix to engage, TonB Q160C did not interact at either a residue central to the mechanically weak FepA domain (D34C) or a residue at its near boundary with the mechanically recalcitrant FepA domain (G54C).

The combination of F202A, W213A mutations on either side of the TonB amphipathic helix substantially inhibited the normal interaction of TonB Q160C with FepA TonB box residue V16C. As such, the double TonB mutation somewhat mimicked the effect of the FepA TonB box deletion that prevented Q160C interactions with FepA D34C and G54C, suggesting that TonB F202A, W213A might have inhibited the amphipathic helix from engaging the FepA TonB box, preventing the FepA cork from fishing for TonB Q160C.

Consistent with our results revealing movement of the FepA cork, Majumdar et al. engineered intra-cork disulfide bonds in FepA, most of which significantly decreased Fe-enterochelin transport. The transport was restored in the presence of reducing agent, indicating that there are required conformational changes within the cork itself (52).

### Comparison to results from *in vivo* photocrosslinking to study FepA dynamics

We previously identified FepA residues that interact with TonB using *in vivo* photocrosslinking by the reagent pBpa inserted at sites of engineered amber substitutions in *fepA* (40). A potential conformational switch signaling to TonB that ligand (enterochelin) is bound was identified. FepA T32pBpa bound TonB in the absence enterochelin whereas FepA A33pBpa bound TonB in its presence. In the current study, both FepA T32C and A33C interactions occurred with both TonB R204C in the amphipathic helix and Q160C without discrimination, perhaps reflecting an average of ligand-bound and unbound states for FepA. FepA S29pBpa, A42CpBpa, S46pBpa, and T51pBpa in the mechanically weak domain did not significantly photocrosslink to TonB, even though these variants all supported ∼ 75% activity. This could be because the interactions are too rapid to capture, whereas in the current study, the disulfide bond formations would potentially have been aided by the DsbA system (64).

Although the effect of a *dsbA* mutation on disulfide formation in this study was not tested, the effect on TonB triplet homodimer formation are informative and suggest that the frustrating answer is: it depends. Plasmids expressing TonB F125C, G186C, F202C, W213C, Y215C, or F230C substitutions were transformed into an isogenic *dsbA* strain, KP1514 [W3110, Δ(*tonB, P14*::*blaM)*, *dsbA*::*kan*], and the degree of disulfide-linked triplet homodimer formation was compared to previous results from a *dsbA+* strain (27). TonB F125C, TonB Y215C and TonB F230C showed greatly diminished triplet dimer formation in the absence of DsbA, with an intermediate decrease for W213C. In no case was triplet dimer formation entirely abolished. For TonB F202C and TonB G186C, there was no effect of the *dsbA* mutation (Spicer and Postle, unpublished data).

In contrast to the present study, FepA E120pBpa and I145pBpa on the periplasmic face of the FepA cork, did photocrosslink to TonB (40). We do not have an explanation for these differences but note that the techniques are dissimilar, and the disulfide crosslinking studies here were congruent with our studies of *in vivo* FepA cork accessibility (12).

### Current *in vivo* approaches are not amenable to discovery of TonB aromatic residue contacts with FepA

This study revealed the importance of the TonB amphipathic helix core (residues 204-212) in contacting the TonB box of FepA. However, individual Cys substitutions within that core have no phenotype (27). Similarly, the sequences of TBDT TonB boxes contain little residue-specific information and indeed can be swapped for one another (15, 16). It is therefore unlikely that this set of interactions is responsible for the idiosyncratic phenotypes observed for Cys and Ala substitutions of the aromatic residues that boundary the amphipathic helix core—F202 and W213 among others.

We have been, unfortunately, thwarted in our ability to define the sites on FepA where TonB F202 and W213 interact due to two factors, both based on signal-to-noise ratios. First, Cys substitutions at these aromatic residues form a sufficiently high abundance of disulfide-linked triplet homodimers *in vivo* that any side reactions would be swamped out and difficult to interpret (17, 27). Second, there appears to be a region of TonB that cannot be analyzed by in vivo photocrosslinking. We previously used targeted amber mutations in *fepA* and in *exbD* to guide insertion of the photocrosslinkable amino acid pBpa and generate crosslinks at unknown sites in TonB *in vivo*. In those studies, both the *fepA* and *exbD* amber mutations fully incorporate the pBpa and result in full-length proteins (40, 65).

We were hopeful that a reciprocal approach using targeted amber mutations in *tonB* at the carboxy terminal aromatic residues would be fruitful, however, we were thwarted by failure to incorporate sufficient pBpa, except small amounts and only after highest overexpression, such that ∼ 85-100% of the TonB was present as the truncated amber mutant form or its degradation product. Given the dominant negative gene dosage effect of *tonB* overexpression, the high level of incomplete TonB fragments would have obscured meaningful interpretations (3, 66). There may be something unusual about this particular region of *tonB* during translation since we have been able to successfully incorporate pBpa into *tonB* at engineered amber sites in the transmembrane domain (Postle and Guzek, unpublished data).

### Surveillance of FepA periplasmic β-strand turns

TonB binds to transporters whether or not it is “energized”, although it is still not clear what that term means mechanistically. For example, inactive TonB H20A formaldehyde-crosslinks to FepA *in vivo* (18) and purified inactive carboxy terminal domains of TonB bind with varying affinities to purified transporters *in vitro* (19, 21, 22). Here we identified the first *in vivo* interactions between the TonB amphipathic helix and the FepA β-strand turns, the majority of which appeared to represent interactions with inactive TonB.

The periplasmic β-strand turns are candidates for one or more TonB-FepA binding sites since they protrude more deeply into the periplasm than the face of the cork does (Fig. 11). We chose Cys substitutions at Asp or Glu residues (and Pro243) in the periplasmic β-strand turns as most likely to be required for FepA activity and were surprised they were all functional. The fact that they had little to no effect on activity, builds on and confirms a tolerance to mutation that generally characterizes TBDTs (36, 40), where only certain structurally disruptive mutations such as a Leu-to-Pro mutation in the TonB box, its complete deletion, or Arg-to-Pro in the lock region residue 75 lead to TBDT inactivation (4, 15, 55).

Interaction of TonB N208C with the FepA periplasmic β-strand turn 5 D422C variant was striking for two reasons: first, because it was so abundant--as abundant as the control TonB Q160C-FepA V16C interaction, and second because it was impervious to the presence of the inactivating TonB H20A substitution. In contrast, TonB R212C formed complexes with Cys residues in several β-strand turns, the majority of which decreased if TonB was inactivated by the H20A mutation.

Phage panning using a purified TonB carboxy terminus identifies TonB interaction sites on FhuA sites corresponding to barrel turns 1 and 2, represented in this study by FepA D185C and P243C (56). We did not observe interaction of TonB with D185, but we did observe R204C, N208C and R212C interacting with P243C; interactions by N208C and R212C were both H20-dependent.

Considering all the results, this study demonstrated that the hydrophilic face of the essential TonB amphipathic helix was used for contacts throughout FepA. It also validated the idea that, *in vivo*, there are certain TonB-FepA contacts made by active TonB, with a different set of contacts that do not lead to energy transduction events. The contacts made by inactive “unenergized” TonB, some of which were quite abundant, could be consistent with membrane surveillance and conformational sampling (27, 46) where TonB discriminates between a TBDT and a porin, or searches for a ligand-loaded TBDT (67). Because the ligand enterochelin was present throughout the experiments here, it will be important to determine which TonB-FepA interaction sites are ligand-dependent. If any *in vivo* TonB-FepA interactions are H20-dependent as well as ligand-dependent, they would constitute candidates important for energy transduction.

## METHODS AND MATERIALS

### Bacterial strains & plasmids

The strains and plasmids used in this study are listed in Table 2. All bacteria are derivatives of *Escherichia coli* K-12 strain W3110. KP1491 was constructed by P1_vir_ transduction of the Δ(*tonB, P14)::blaM* cassette from KP1477 into KP1489 (W3110 Δ*fepA*).

**Table 2:**
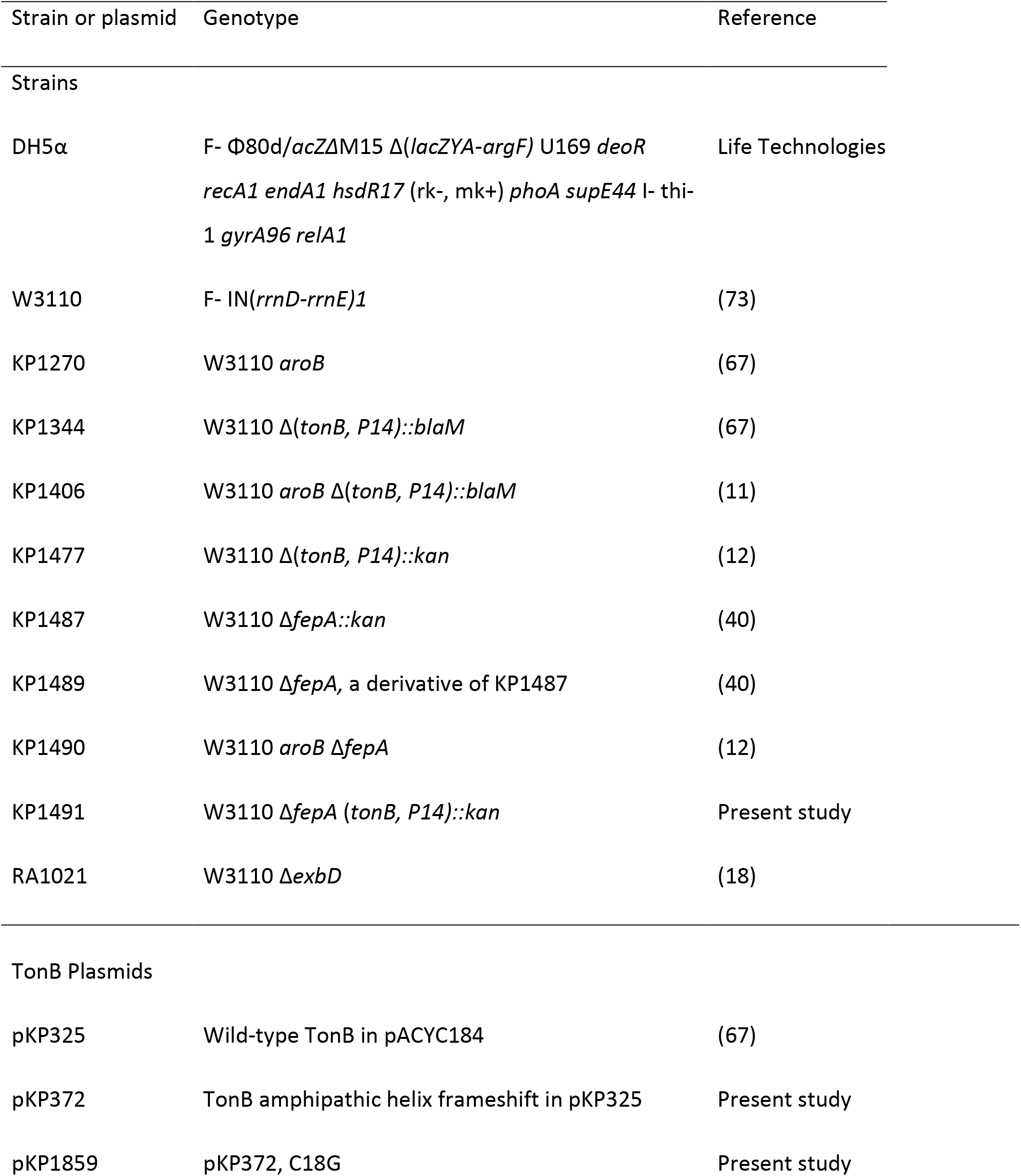
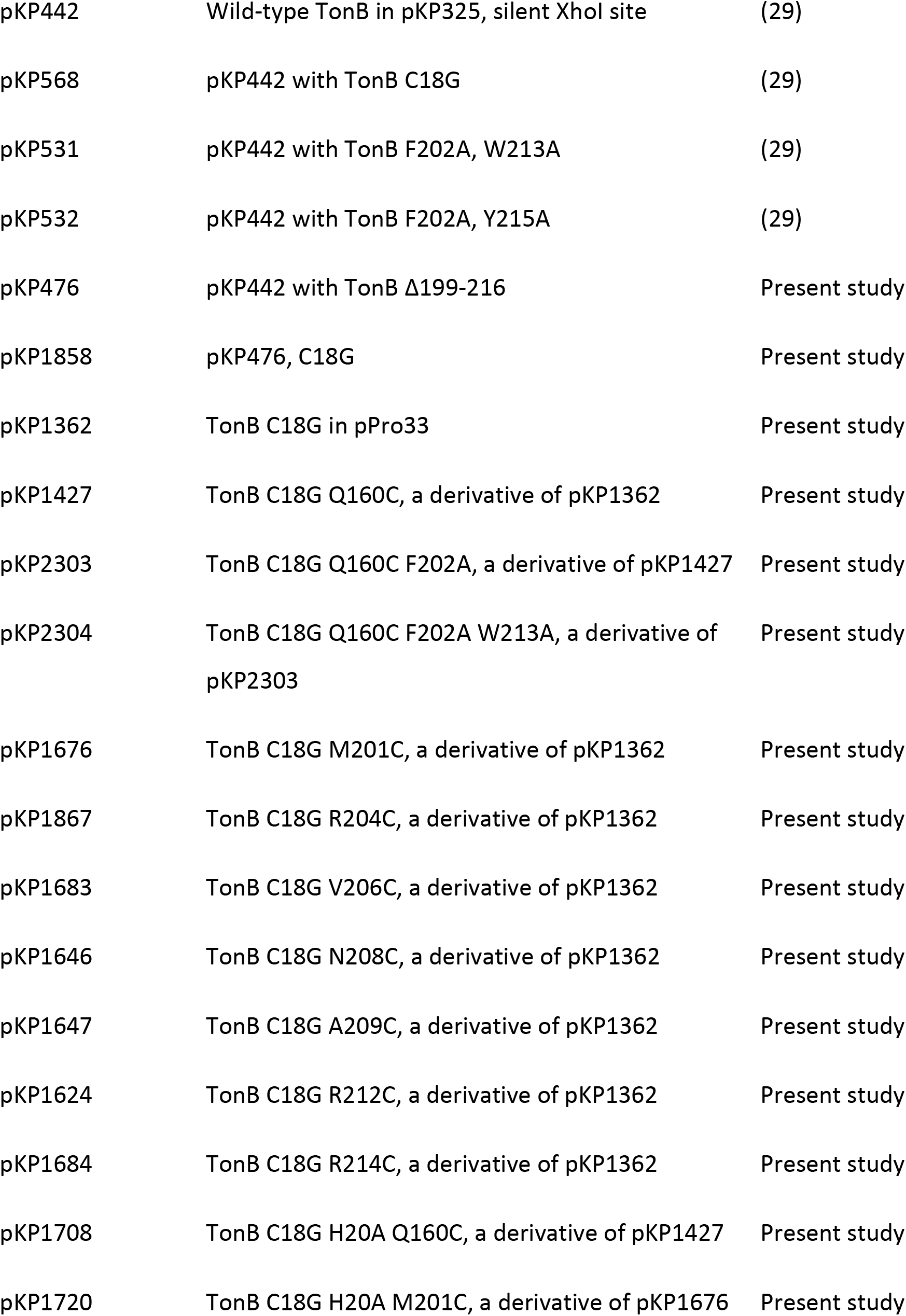
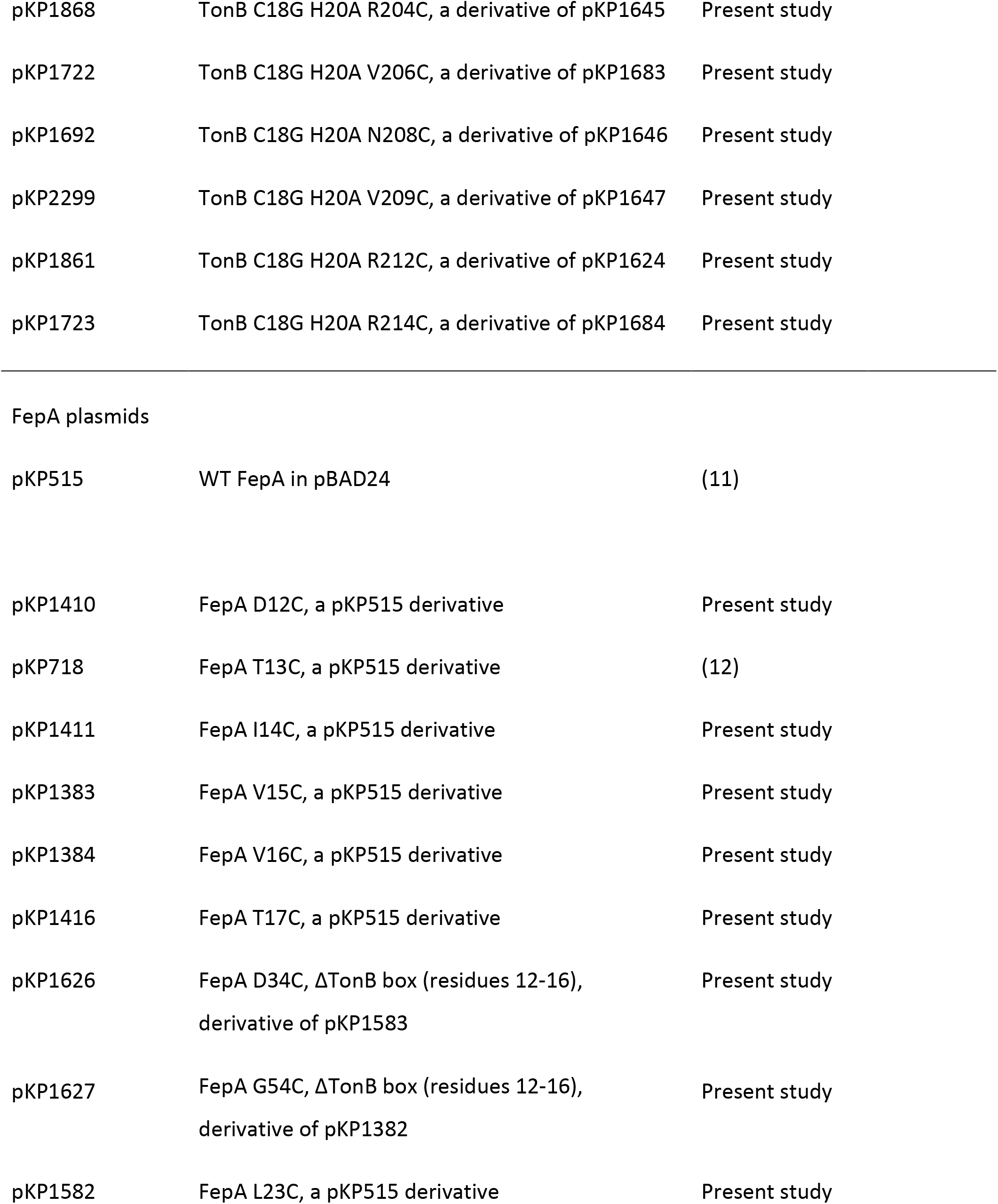
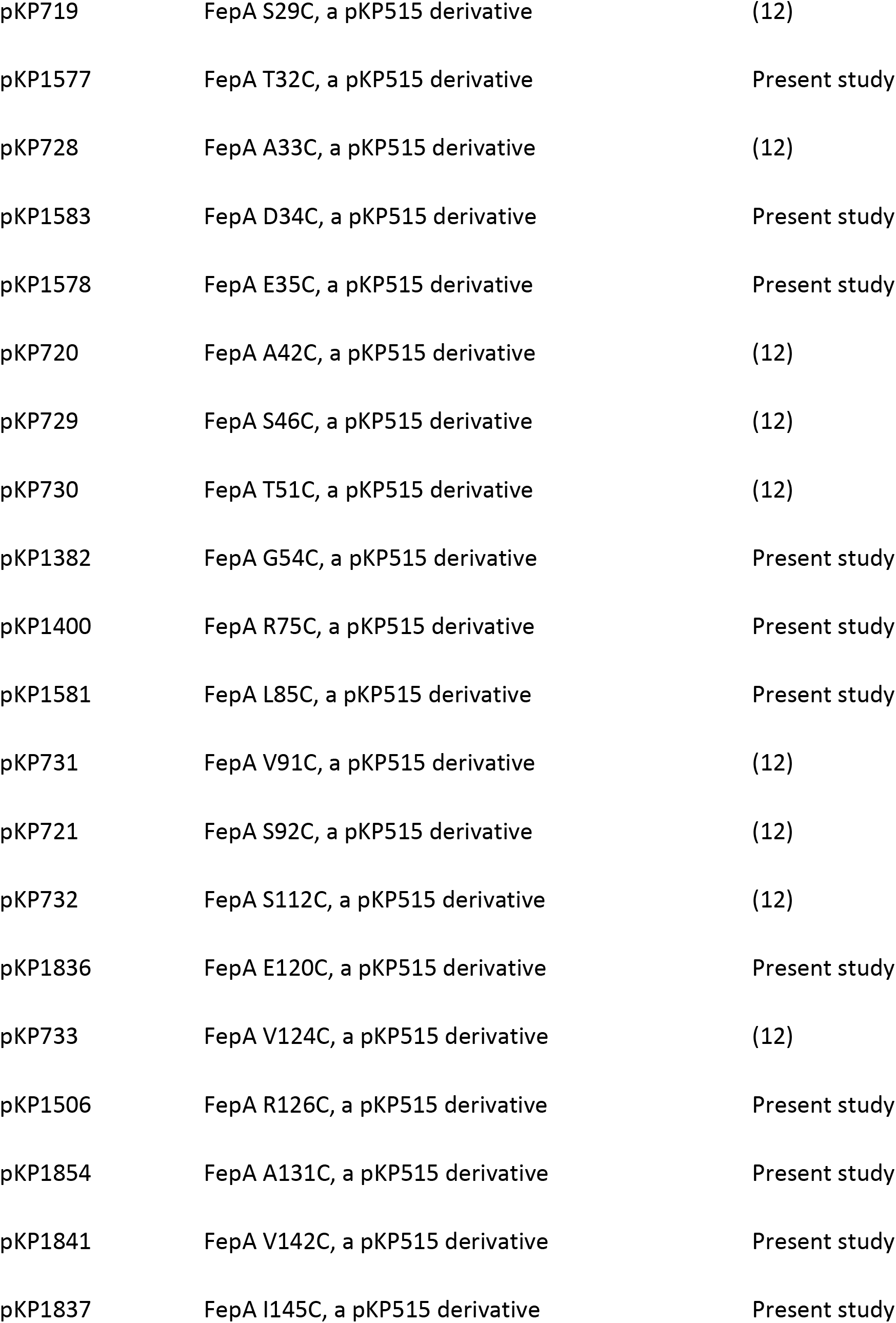
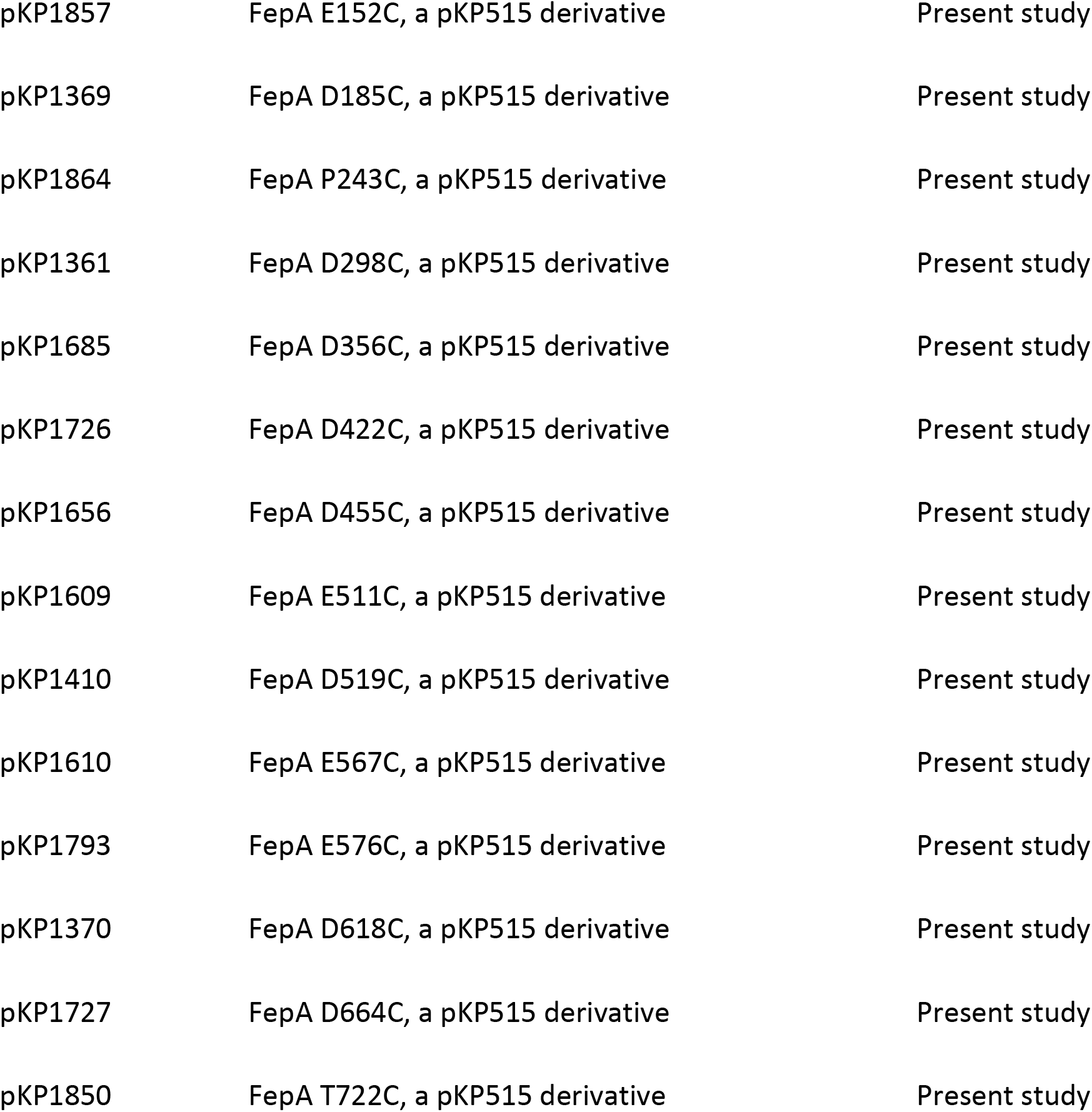
Strains and plasmids

| Strain or plasmid | Genotype | Reference |
| --- | --- | --- |
| Strains |  |  |
| DH5α | F- Φ80d/*acZΔM15 Δ(lacZYA-argF)* U169 *deoR*<br>*recA1 endA1 hsdR17* (rk-, mk+) *phoA supE44* l- thi-<br>1 *gyrA96 relA1* | Life Technologies |
| W3110 | F- IN(*rrnD-rrnE*)1 | (73) |
| KP1270 | W3110 *aroB* | (67) |
| KP1344 | W3110 Δ(*tonB, P14*)::*blaM* | (67) |
| KP1406 | W3110 *aroB* Δ(*tonB, P14*)::*blaM* | (11) |
| KP1477 | W3110 Δ(*tonB, P14*)::*kan* | (12) |
| KP1487 | W3110 Δ*fepA*::*kan* | (40) |
| KP1489 | W3110 Δ*fepA*, a derivative of KP1487 | (40) |
| KP1490 | W3110 *aroB* Δ*fepA* | (12) |
| KP1491 | W3110 Δ*fepA* (*tonB, P14*)::*kan* | Present study |
| RA1021 | W3110 Δ*exbD* | (18) |
| TonB Plasmids |  |  |
| pKP325 | Wild-type TonB in pACYC184 | (67) |
| pKP372 | TonB amphipathic helix frameshift in pKP325 | Present study |
| pKP1859 | pKP372, C18G | Present study |
| pKP442 | Wild-type TonB in pKP325, silent XhoI site | (29) |
| pKP568 | pKP442 with TonB C18G | (29) |
| pKP531 | pKP442 with TonB F202A, W213A | (29) |
| pKP532 | pKP442 with TonB F202A, Y215A | (29) |
| pKP476 | pKP442 with TonB Δ199-216 | Present study |
| pKP1858 | pKP476, C18G | Present study |
| pKP1362 | TonB C18G in pPro33 | Present study |
| pKP1427 | TonB C18G Q160C, a derivative of pKP1362 | Present study |
| pKP2303 | TonB C18G Q160C F202A, a derivative of pKP1427 | Present study |
| pKP2304 | TonB C18G Q160C F202A W213A, a derivative of pKP2303 | Present study |
| pKP1676 | TonB C18G M201C, a derivative of pKP1362 | Present study |
| pKP1867 | TonB C18G R204C, a derivative of pKP1362 | Present study |
| pKP1683 | TonB C18G V206C, a derivative of pKP1362 | Present study |
| pKP1646 | TonB C18G N208C, a derivative of pKP1362 | Present study |
| pKP1647 | TonB C18G A209C, a derivative of pKP1362 | Present study |
| pKP1624 | TonB C18G R212C, a derivative of pKP1362 | Present study |
| pKP1684 | TonB C18G R214C, a derivative of pKP1362 | Present study |
| pKP1708 | TonB C18G H20A Q160C, a derivative of pKP1427 | Present study |
| pKP1720 | TonB C18G H20A M201C, a derivative of pKP1676 | Present study |
| pKP1868 | TonB C18G H20A R204C, a derivative of pKP1645 | Present study |
| pKP1722 | TonB C18G H20A V206C, a derivative of pKP1683 | Present study |
| pKP1692 | TonB C18G H20A N208C, a derivative of pKP1646 | Present study |
| pKP2299 | TonB C18G H20A V209C, a derivative of pKP1647 | Present study |
| pKP1861 | TonB C18G H20A R212C, a derivative of pKP1624 | Present study |
| pKP1723 | TonB C18G H20A R214C, a derivative of pKP1684 | Present study |
| FepA plasmids |  |  |
| pKP515 | WT FepA in pBAD24 | (11) |
| pKP1410 | FepA D12C, a pKP515 derivative | Present study |
| pKP718 | FepA T13C, a pKP515 derivative | (12) |
| pKP1411 | FepA I14C, a pKP515 derivative | Present study |
| pKP1383 | FepA V15C, a pKP515 derivative | Present study |
| pKP1384 | FepA V16C, a pKP515 derivative | Present study |
| pKP1416 | FepA T17C, a pKP515 derivative | Present study |
| pKP1626 | FepA D34C, ΔTonB box (residues 12-16),<br>derivative of pKP1583 | Present study |
| pKP1627 | FepA G54C, ΔTonB box (residues 12-16),<br>derivative of pKP1382 | Present study |
| pKP1582 | FepA L23C, a pKP515 derivative | Present study |
| pKP719 | FepA S29C, a pKP515 derivative | (12) |
| pKP1577 | FepA T32C, a pKP515 derivative | Present study |
| pKP728 | FepA A33C, a pKP515 derivative | (12) |
| pKP1583 | FepA D34C, a pKP515 derivative | Present study |
| pKP1578 | FepA E35C, a pKP515 derivative | Present study |
| pKP720 | FepA A42C, a pKP515 derivative | (12) |
| pKP729 | FepA S46C, a pKP515 derivative | (12) |
| pKP730 | FepA T51C, a pKP515 derivative | (12) |
| pKP1382 | FepA G54C, a pKP515 derivative | Present study |
| pKP1400 | FepA R75C, a pKP515 derivative | Present study |
| pKP1581 | FepA L85C, a pKP515 derivative | Present study |
| pKP731 | FepA V91C, a pKP515 derivative | (12) |
| pKP721 | FepA S92C, a pKP515 derivative | (12) |
| pKP732 | FepA S112C, a pKP515 derivative | (12) |
| pKP1836 | FepA E120C, a pKP515 derivative | Present study |
| pKP733 | FepA V124C, a pKP515 derivative | (12) |
| pKP1506 | FepA R126C, a pKP515 derivative | Present study |
| pKP1854 | FepA A131C, a pKP515 derivative | Present study |
| pKP1841 | FepA V142C, a pKP515 derivative | Present study |
| pKP1837 | FepA I145C, a pKP515 derivative | Present study |
| pKP1857 | FepA E152C, a pKP515 derivative | Present study |
| pKP1369 | FepA D185C, a pKP515 derivative | Present study |
| pKP1864 | FepA P243C, a pKP515 derivative | Present study |
| pKP1361 | FepA D298C, a pKP515 derivative | Present study |
| pKP1685 | FepA D356C, a pKP515 derivative | Present study |
| pKP1726 | FepA D422C, a pKP515 derivative | Present study |
| pKP1656 | FepA D455C, a pKP515 derivative | Present study |
| pKP1609 | FepA E511C, a pKP515 derivative | Present study |
| pKP1410 | FepA D519C, a pKP515 derivative | Present study |
| pKP1610 | FepA E567C, a pKP515 derivative | Present study |
| pKP1793 | FepA E576C, a pKP515 derivative | Present study |
| pKP1370 | FepA D618C, a pKP515 derivative | Present study |
| pKP1727 | FepA D664C, a pKP515 derivative | Present study |
| pKP1850 | FepA T722C, a pKP515 derivative | Present study |

Plasmids pKP1858 and pKP1859 were created from pKP476 and pKP372 respectively, using polymerase chain reaction (PCR)-based site-directed mutagenesis as previously described (26) to create C18G substitutions in both plasmids. The majority of plasmids encoding *tonB* mutants were derived from pKP1362 (*tonB* C18G), which was constructed by cloning *tonB* C18G from pKP568 into the Sph1 site of pPro33, allowing for expression from the propionate promoter (68). Plasmids encoding *fepA* mutants were derived from pKP515 where *fepA* is expressed from the arabinose promoter in pBAD24 (11). Mutations were engineered through PCR-based site-directed mutagenesis as previously described (26). The coding region of each engineered mutant gene was confirmed through Sanger sequencing at the Pennsylvania State University Nucleic Acid Facility.

### Growth media and culture conditions

Liquid cultures were grown at 37°C with aeration in LB broth or in M9 minimal salts supplemented with 1% glycerol, 0.2% vitamin-free casamino acids, 40 µg ml^-1^ of tryptophan, 4 µg ml^-1^ of thiamine, 1 mM MgSO_4_ and 0.5 mM CaCl_2_. For disulfide-crosslinking, the M9 minimal salts medium was further supplemented with 10 µM Fe (as ferric chloride). For [^55^Fe]-enterochelin transport assays, the M9 minimal salts medium was supplemented with 1.85 µM Fe (as ferric chloride) as well as 40 µg ml^-1^ of tyrosine and 40 µg ml^-1^ of phenylalanine to facilitate growth of *aroB* strains on which they were performed (69). Chloramphenicol at 34 µg ml^-1^ and ampicillin at 100 µg ml^-1^ were used to maintain TonB and FepA plasmids respectively. The TonB plasmids were induced with the following concentrations with sodium propionate to achieve chromosomal levels; TonB C18G (10 mM), TonB C18G M201C (10 mM), TonB C18G R204C (0.5 mM), TonB C18G V206C (10 mM), TonB C18G N208C (10 mM), TonB C18G A209C (10 mM), TonB C18G R212C (1 mM), TonB C18G R214C (10 mM), all TonB C18G H20A cysteine substitutions (15 mM). FepA cysteine substitutions were not induced as the base expression level approximated chromosomally encoded FepA levels in cells grown with 1.85 µM Fe.

### Sucrose density gradient fractionations

Sucrose density gradient fractionation was carried out essentially as described previously (43) with some modifications. Strain KP1344 containing plasmids pKP1858 or pKP1859 was grown as described above, in the presence of 0.002% arabinose and 0.1% arabinose respectively, to mid exponential phase. Cells were harvested and lysed by French pressure cell at 4° C. The cell lysate supernatant was applied to the top of the sucrose gradient and centrifuged in a Beckman SW40 rotor at 35,000 r.p.m. for 19 hours at 4° C. Collected fractions were precipitated with an equivalent volume of 20% trichloroacetic acid (TCA) at 4° C, and suspended in Laemmli sample buffer (70) (LSB) at 95°C for 5 minutes. 10 µl of each sample was electrophoresed on 12% SDS-polyacrylamide gels and then immunoblotted with TonB 4F1 monoclonal antibodies (71).

### In vivo formaldehyde crosslinking

Strains were subcultured 1:100 from saturated LB cultures into supplemented M9 minimal salts medium supplemented with L-arabinose as described in the Figure 17 legend and 34 µg ml^-1^ chloramphenicol without added iron. Cells were harvested at an A_550_ of 0.5 in 1 ml aliquots, centrifuged and aspirated. The pellet was suspended in 938 µl of 100 mM sodium phosphate buffer at pH 6.8 to which 62.5 µl of 16% formaldehyde was added and incubated for 15 minutes at 22°C. Cells were then pelleted, suspended in 50 µl of 2x LSB (twice the usual concentration) and heated for 5 minutes at 60°C. The samples were electrophoresed on 11% SDS-polyacrylamide gels and then immunoblotted with TonB 4F1 monoclonal antibodies or anti-ExbD polyclonal antibodies (32, 71).

### In vivo disulfide crosslinking

KP1491 harboring pairwise combinations of plasmid-encoded TonB and plasmid encoded FepA were subcultured 1:100 from saturated LB cultures into supplemented M9 minimal salts medium and grown with appropriate antibiotics to A_550_ = 0.45. 0.4 OD ml^-1^ of cells were harvested by centrifugation and precipitated with an equal volume of 4°C 20% TCA to stop the proteolysis of TonB that occurs in LSB at 95°C when TCA is not used (72). The TCA-precipitated pellets were boiled at 95°C for 10 minutes in 100 µl of LSB with 50 mM iodoacetamide to block any remaining free cysteines to prevent *in vitro* disulfide crosslinking. All samples were analyzed on 9% SDS-PAGE gels and followed by immunoblot analysis with TonB 4F1 monoclonal antibody and FepA polyclonal antibodies (32).

To eliminate the possibility that disulfide crosslinks formed due to the presence of TCA during cell harvesting, the efficiency of crosslinking was compared with and without TCA precipitation. Upon harvesting cells were pelleted without TCA. The cell pellets were boiled at 95°C for 10 minutes in 100 µl of LSB with 50 mM iodoacetamide. TCA slightly enhanced the recovery of both crosslinked complexes, which also still formed in the absence of TCA and the monomer such that levels of crosslinking were proportional to the controls with and without TCA (data not shown).

### [*^55^Fe*]-enterochelin transport

TonB and FepA with individual Cys substitutions were assessed for their initial rates of enterochelin (Sigma-Aldrich) transport as described previously (40). FepA constructs were assayed in KP1490 (W3110 *aroB* Δ*fepA*) whereas TonB constructs were assayed in strain KP1406 (W3110 *aroB* Δ(*tonB, P14)::blaM*). Enterochelin is the sole siderophore synthesized and secreted by *E. coli* K12. The *aroB* mutation prevents enterochelin synthesis and the synthesis of any intermediates that could interfere in the assay (69).

## ACKNOWLEDGEMENTS

We thank Ryan Guzek for analysis of the TonB pBpa substitutions and construction of pKP1836 and pKP1837; Bradley Spicer for construction of pKP1382, pKP1400, pKP1403, pKP1427, FepA Cys substitutions in the TonB box, and analyzing effects of a *dsbA* strain on TonB triplet homodimer formation; Suzanne Wardell for construction of pKP372; Shaima El Mowafi for construction of pKP1369 and pKP1370; Siti Kamarudin for construction of pKP1858, pKP1859, and pKP1861; Surendran Devanathan for construction of KP1491; Cheryl Swayne for construction of pKP1362; Raka Ghosh for construction of pKP1581; Yu-An Liu for construction of pKP1361; Larissa Goldman for construction of pKP1506; Gui Teng Chua for construction of pKP1841, pKP1850, and pKP1854; Elizabeth McFadden for the formaldehyde crosslinking experiment on the TonB double Ala variants, and Glenn Hwang for construction of pKP2299, pKP2303, pKP2304, the analysis of amphipathic helix Cys substitutions interactions with the FepA TonB box and excellent technical assistance. We thank the *E. coli* Genetic Stock Center and the Jonathan Beckwith lab for their *dsbA*::kan strain, RI90. Support from NIGMS grant GM112710 is gratefully acknowledged.

